# The linker histone H1.4 condenses chromatin in maturing postmitotic neurons

**DOI:** 10.64898/2026.07.28.741367

**Authors:** Andrew I. Aldridge, Martine W. Tremblay, Vijyendra Ramesh, Xiaolin Wei, Yong-Hui Jiang, Anne E. West

**Affiliations:** Department of Neurobiology, Duke University School of Medicine, Durham, NC 27710; Department of Cell Biology, Duke University School of Medicine, Durham, NC 27710; Department of Genetics, Neuroscience, and Pediatrics, Yale University School of Medicine, New Haven, CT 06510

## Abstract

H1 histones are abundant nuclear proteins that bind to linker DNA at the entry and exit points of nucleosomes. Although individual H1 family members play partially redundant roles in chromatin organization, the discovery of disease-associated mutations in H1 genes has raised interest in their cell-type specific functions. Heterozygous, de novo frameshift mutations in *H1-4* cause the neurodevelopmental disorder Rahman Syndrome, which is characterized by mild to severe intellectual disability along with other neurological and morphological features. H1.4 has mostly been studied in dividing cells, and its expression in the brain was poorly understood. Here we characterized the expression of *H1f4* mRNA and H1.4 protein in the brains of male and female mice across postnatal development. Using an epitope-tagged *H1f4* knockin mouse, we show that this linker histone is robustly expressed throughout the brain including in mature, post-mitotic neurons of adult mice. By chromatin immunoprecipitation, we observe that H1.4 binds broadly across the genome in neural progenitors and accumulates in heterochromatin over the course of neuronal maturation. Finally we show that developmental maturation of chromatin compaction in cerebellar granule neurons is disrupted in *H1f2/H1f4* double knockout mice. These data raise the possibility that the neurological changes in Rahman Syndrome may arise from disrupted functions of histone H1.4 in neurons.

## INTRODUCTION

The H1 family of linker histones play fundamental structural organizing roles in chromatin. H1 histones bind to linker DNA at the entry and exit sites of nucleosomes, and they act to compact chromatin into higher order structures (1). There are 11 separate genes encoding histone H1s in humans and mice, with 7 somatic H1s (*H1f0-H1f5*, and *H1f10* in mice) and 4 that are germline specific. Only two of the somatic H1 histones (*H1f0* and *H1f10*) are polyadenylated, and these have been characterized as replacement histones that are incorporated into chromatin independently of the cell cycle. By contrast the other five somatic H1s (*H1f1-H1f5*) generally lack polyA tails and their mRNAs are most highly expressed in dividing cells during S phase (2,3). This has led to their designation as “replication-dependent” H1s, though there has long been some evidence that this group of H1s might also have functions in differentiated cells (2,4,5).

Knockout of any one of the somatic H1s in mice, including the replacement histone *H1f0,* has minimal effects on development or organismal survival (6,7). By contrast triple knockout of *H1f2, H1f3,* and *H1f4* lowers total histone H1 levels to 50% of normal and is lethal at mid-gestation (8). These data indicate that compensation between members of the H1 family is essential for organismal survival, but they offer little insight into specific functions of the H1s in different tissues. More recently, missense mutations in H1s have been identified that are associated with diseases of particular cell types. For example, recurrent missense mutations of *H1-2* and *H1-4* have been identified in B-cell lymphomas. In *H1f2/H1f4* double KO mice, germinal center B-cells displayed large scale genomic decompaction and increased expression of stem-cell genes, revealing the importance of these two H1 histones for chromatin organization in B-cells (9).

Rahman Syndrome is a rare neurodevelopmental disorder caused by de novo, heterozygous frameshift mutations that disrupt the C-terminal domain of the gene encoding histone H1.4 (10). These frameshift mutations result in expression of an H1.4 protein that lacks the highly negatively charged C-terminus, which is thought to contribute to higher order chromatin condensation, and gains a novel C-terminal domain that may have neomorphic consequences (10–13). We previously showed that expressing the frameshift mutant H1.4 in rat hippocampal neurons disrupts neuronal gene expression and impairs neuronal firing (14). These data suggest that Rahman Syndrome mutations could act in neurons to disrupt brain function. However, when and where histone H1.4 is expressed in the brain and what role it plays for chromatin organization in postmitotic cells was poorly understood.

Here to determine the expression pattern and function of H1.4 in the mouse brain, we generated a 3xFLAG-Myc epitope-tagged *H1f4* knockin mouse. Combining western blotting and immunohistochemistry with RT-PCR and fluorescence in situ, we demonstrate that although *H1f4* mRNA expression declines when neurons leave the cell cycle, H1.4 protein continues to accumulate in most cell types throughout the brain into adulthood. Our ChIP-seq data show that H1.4 accumulates in heterochromatin over time as neurons mature and we conducted chromatin conformation studies that show how double knockout of *H1f2* and *H1f4* disrupts chromatin organization in adult cerebellar granule neurons. These data demonstrate that histone H1.4 is highly expressed and functionally important for the organization of chromatin in postmitotic cells neurons, and they raise the possibility that aberrant function of H1.4 in these neurons could contribute to intellectual disability in Rahman Syndrome.

## MATERIALS AND METHODS

### Mouse strains

*H1f2/4* double knockout (dKO) germline mice were a gift from the laboratory of Arthur Skoultchi (7).C57BL/6J mice were obtained from The Jackson Laboratory (#000664; RRID:IMSR_Jax:000664) and CD-1 IGS mice (#Strain 022, RRID:IMSR_CRL:022) were obtained from Charles River Laboratories. 3xFLAG-Myc*-H1f4* mice were created at Duke University by the Rodent Cancer Models Shared Resource. Briefly, a homology independent target insertion (HITI)-based targeting vector approach with neomycin (neo) selection was used (15). Knock-in targeting and mock control vectors were sequence verified, linearized, and a PCR assay was validated for screening targeted ES cell clones. The G4 129/B6N hybrid ES line were obtained by the Duke Transgenic core by MTA from Mount Sinai Hospital in Toronto (16). G4 ES cells of positive clones were microinjected into ICR morulae to produce chimeric mice. The following 3xFLAG-Myc sequence was inserted immediately after the endogenous *H1-4* start codon: GACTACAAAGACCATGACGGTGATTATAAAGATCATGATATCGATTACAAGGATGAC GATGACAAGGGAGGCGAACAAAAGTTGATTTCTGAAGAAGATTTG. Positive mice were then bred to B6.Cg-Tg(ACTFLPe)9205Dym/J (Jackson Labs #005703) mice expressing Flp recombinase to remove the FRT-flanked neomycin cassette. All 3xFLAG-Myc-*H1f4* mice used in this study were heterozygous for the modified allele. Both male and female mice were used for all studies in this project. All experiments were conducted under a protocol approved by the Duke University Institutional Animal Care and Use Committee.

### RT-qPCR

RNA was isolated from mouse brain tissue using TRIzol Reagent (Thermo cat #15596026). RNA was treated with DNAse (NEB cat #M0303) prior to cDNA synthesis (Invitrogen cat #18080051) with random hexamers or oligo dT only if specifically indicated in the text per manufacturers protocols. qPCR was performed using an Applied Biosystems Quantstudio 3 Thermal cycler with Power SYBR Green PCR Master Mix (Thermo cat #4367659) and PCR primers targeting the gene of interest (Table 1).

**Table 1:**
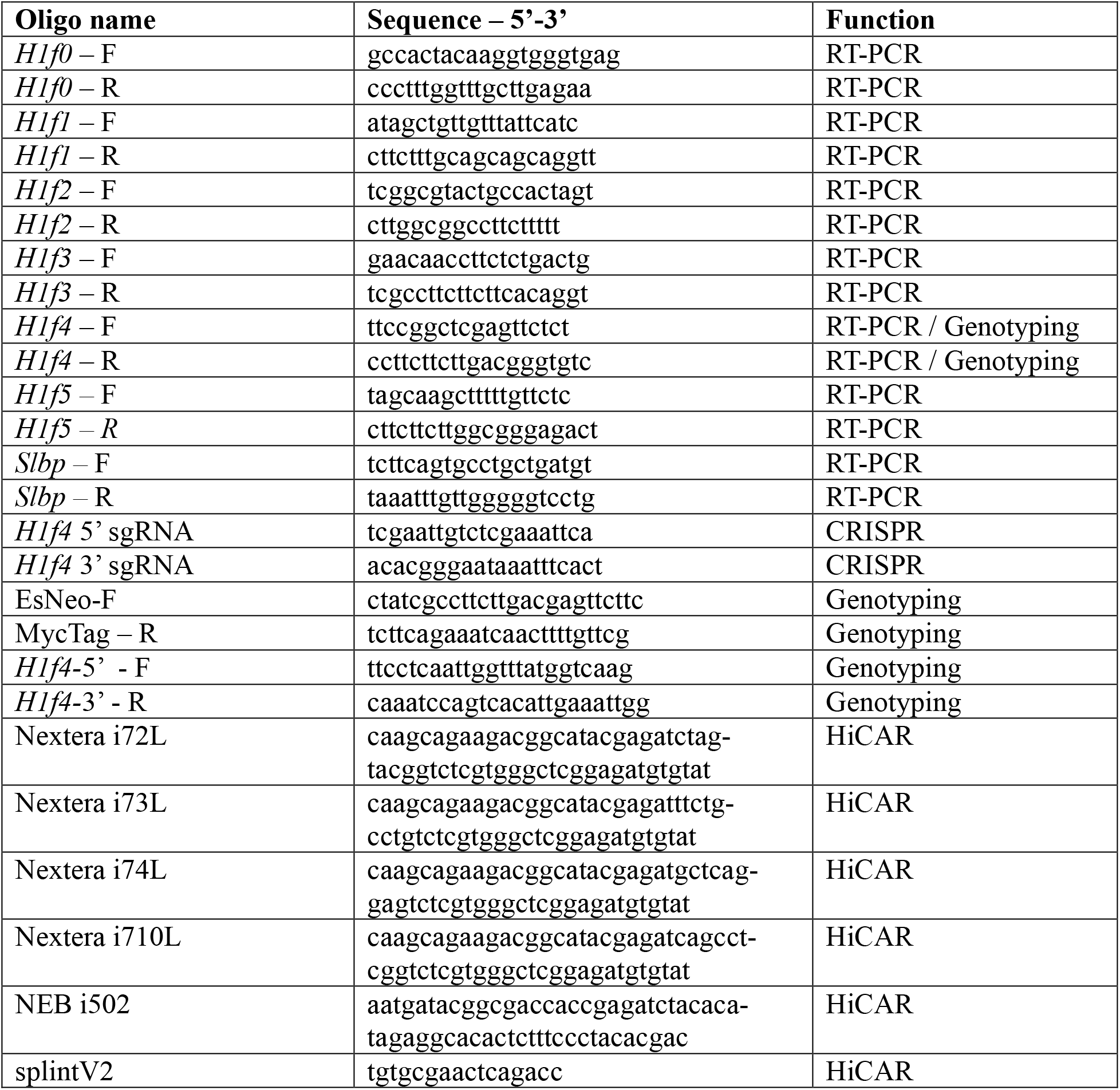
PCR primers used in this study.

### In-situ hybridization

In situ hybridization was performed using ACD RNAScope Fluorescent Multiplex kit on fresh-frozen brains of WT animals. Specific additions/modifications to the protocol are as follows: slice sections at 20µm, protease IV antigen retrieval (ACD cat #322340), and AMP 4-FL-Amp-A. Probes used for this study are: *H1f2* (ACD cat #584831-C2), *H1f3* (ACD cat #584841-C2), *H1f4* (ACD cat #584821-C2), and *Map2* (ACD cat #431151). Images were gathered on a Leica SP8 confocal microscope and processed in ImageJ.

### Western blots

Histones were extracted from mouse brain tissue using acid extraction. Briefly, nuclei were isolated and treated with 0.2M ice-cold hydrochloric acid for 5 minutes. Samples were spun at 16,000g 4°C for 5 minutes. To precipitate histones, one quarter volume of 100% (w/v) trichloroacetic acid was added to the supernatant and incubated for 5 minutes on ice. Samples were spun at 16,000g 4°C for 5 minutes. Isolated histones were then washed twice with 100µL ice cold acetone. Histone extracts were boiled in Laemmli buffer for 10 minutes and run on a 15% SDS-PAGE gel and transferred to a nitrocellulose membrane. Membranes were blocked for 1 hour RT in 5% milk(w/v) in TBST, probed with primary antibody (rabbit anti-FLAG, Sigma-Aldrich cat #F7425, RRID:AB_439687, 1:1000; rabbit anti-H1.0, Abcam cat #ab125027, RRID:AB_11000797, 1:1000; and mouse anti-H3, Millipore cat #05-499, RRID:AB_309763, 1:1,000) overnight in blocking solution, rinsed, and incubated with fluorescent conjugated secondary antibodies for 1 hour at RT (Biotium cat #20078-1, RRID:AB_10852687, 1:5000, Biotium cat #20067-1, RRID:AB_10871686, 1: 5000). Blots were imaged on a Licor Odyssey and quantified using ImageJ. Statistical comparisons were conducted using Graphpad Prism one-way ANOVA for time.

### Immunofluorescence

Animals were perfused using PBS followed by 4% PFA. Tissue was post-fixed overnight in 4% PFA, then cryoprotected with 30% sucrose solution in PBS. Brains were cut into 40µM sections on a freezing microtome. Sections were blocked and permeabilized for 1 hour RT in PBS+1% Triton-X and 10% Donkey serum prior to incubation overnight at 4°C with primary antibody: goat anti-Myc, Abcam cat #ab9132, RRID:AB_307033, 1:500; mouse anti-calbindin Sigma-Aldrich cat #C9848, RRID:AB_476894 1:1000; rabbit anti-MeCP2, EnCor Biotechnology cat #RPCA-MeCP2, RRID:AB_2572345, 1:1000; rabbit anti-IBA1, FUJIFILM Wako Pure Chemical Corporation cat #019-19741, RRID:AB_839504 1:1000; mouse anti-GFAP, Sigma-Aldrich cat #G3893, RRID:AB_477010 1:100; anti-Zic 1/2 C terminus, courtesy of R. Segal, Harvard Medical School(17). Sections were then washed 3×10 minutes in PBS prior to secondary antibody incubation for 1 hour RT, 1:1000 dilution: donkey anti-goat 488, Thermo Fisher Scientific cat #A-11055, RRID:AB_2534102; donkey anti-rabbit 647, Thermo Fisher Scientific cat #A-31573, RRID:AB_2536183, donkey anti-mouse 594, Thermo Fisher Scientific cat #A-21203, RRID:AB_2535789; donkey anti-chicken 594, Thermo Fisher Scientific cat #A78951, RRID:AB_2921073. Sections were washed 3×10 minutes in PBS and then incubated in Hoechst for 5 minutes RT. Sections were mounted using Prolong Gold antifade reagent (Thermo Fisher cat #P36930). Slides were imaged on a Zeiss 780 inverted confocal microscope. Images were processed with ImageJ. In specific cases antigen retrieval was performed to maximize nuclear signal. Immunostaining was done as described, but prior to permeabilization sections were submerged in 10mM sodium citrate (pH 6.0) at 95°C for 10 minutes. Sections were allowed to cool in the buffer and then washed twice in PBS before continuing.

### Chromatin immunoprecipitation

Cerebellum from each of three P60 3xFLAG-Myc*-H1f4* mice were used for ChIP. For GNPs, cerebella of 3-5 3xFLAG-Myc*-H1f4* mice were dissected at P7. Cells were dissociated and separated on a Percoll gradient as previously described (18). 30-40 million GNPs were used per replicate. For adult cerebellum, tissue was dounced in 1mL of 0.5% formaldehyde (w/v) in PBS and rotated for 10 minutes RT. Formaldehyde was added to GNPs in suspension. Formaldehyde fixation was quenched with 0.2M glycine and rotated an additional 10 minutes RT, followed by two washes with ice cold PBS. Samples were treated with 600ul lysis buffer (1% SDS (w/v), 10mM EDTA, and 50mM Tris, pH 8.1) and sonicated with a Bioruptor Pico (Diagenode), with 30 seconds on/off cycles to an average size range of 150–350bp as determined by agarose gel electrophoresis. Sonicated chromatin was diluted 10-fold in dilution buffer (0.01% SDS, 1.1% Triton X-100, 1.2mM EDTA, 16.7mM Tris-HCl, pH 8.1, 167mM NaCl). Prior to immunoprecipitation, 6µl of antibody (goat anti-Myc, Abcam cat #ab9132, RRID:AB_307033) was incubated with 100µl of Dynabeads Protein G (Invitrogen 10004D) for 4 hours at 4°C. The antibody-bead conjugate was then washed twice with cold PBS, and added to 1ml of cell lysate for overnight immunoprecipitation at 4°C. The following day, bead-bound DNA-protein complexes were washed, eluted, and purified using a PCR purification kit (Qiagen cat #28104). ChIP-seq libraries were made using the NEBNext Ultra II DNA Library Prep Kit for Illumina (New England Biolabs cat #E7645S) and paired-end sequencing was performed on a Novaseq 6000 at the Duke Sequencing and Analysis Core Resource.

### HiCAR

For HiCAR, either isolated GNPs from P7 CD1 mice or adult cerebellum from P60 *H1f2/H1f4* dKO mice and C57/BL/6J mice were used (two each). HiCAR was performed following published protocols(19,20). Briefly, nuclei were fixed in 1% formaldehyde for 10 minutes RT. Formaldehyde fixation was quenched with 0.2M glycine and rotated an additional 10 minutes. 150,000 nuclei per sample were used for each HiCAR experiment. Nuclei and Tn5 transposome were incubated with agitation at 37°C for 1 hour. Tagmented DNA was digested with MseI restriction enzyme (NEB, cat #R0525S) alongside splint2 oligonucleotide (Table 1). In-situ ligation was performed using T4 DNA ligase (NEB, cat #M0202L) for 4 hours RT. Following ligation, samples were reverse-crosslinked overnight at 68°C. DNA was ethanol precipitated at -80°C for 30-60 minutes. DNA was then digested with NIaIII restriction enzyme (NEB, cat #R0125S) at 37°C for 1 hour and cleaned up using 0.9X volume of SPRI beads (Beckman, cat #B23319). DNA concentration was measured using a Qubit instrument and adjusted to a concentration of 1ng/µL. A second round of ligation was performed using T4 DNA ligase incubated for 2 hours RT. DNA was purified using the Zymo DNA clean and concentrator kit (Zymo, cat #D4029) and eluted using 10mM Tris-HCl, pH 8.0. A PCR reaction mixture was set up, using unique Nextera-pcr-i7-primers and NEB i5 primers to amplify and index samples (Table 1). Resulting libraries were checked for fragment size using a bioanalyzer, after which they were pooled and sequenced. 150bp paired-end sequencing was performed at the Duke Sequencing and Analysis Core Resource on a NovaSeq 6000 S-Prime flow cell for GNPs, and a NovaSeq X Plus 10B lane for adult cerebellum.

### RNA-Seq

RNA was harvested from cerebellum from four *H1f2/H1f4* dKO and four C57BL/6J mice. Cerebellar samples were homogenized in Trizol and RNA was isolated using Zymo Direct-zol RNA miniprep kit (Zymo cat #R2052). Sequencing libraries were generated using Watchmaker mRNA Library Prep Kit and sequenced by the Duke Sequencing core on an Illumina NextSeq 1000.

### Computational Methods

#### ChIP

ChIP-Seq reads were processed using the nf-core/chipseq v2.0.0 pipeline. Briefly, reads were quality scored using FastQC v0.11.9, trimmed for adapter sequences using Trimgalore v0.6.7, and aligned to mouse GRCm38 genome using BWA v0.7.17. Due to the low signal-to-noise ratio of Myc-H1.4, immunoprecipitated reads were normalized to input using the bamCompare function of the deepTools2.0 suite. Read counts over chromatin compartments were counted using the featureCounts package v2.0.3. Immunoprecipitated reads were again normalized against input reads.

#### HiCAR

HiCAR reads were processed using the Nextflow jianhong/hicar ver2rc pipeline using default parameters. Briefly, reads were quality scored using FastQC v0.11.9. Reads were mapped to the mouse GRCm38 genome using BWA v0.7.18. Pairs were filtered using Pairtools and quality scored with PairsQC. ATAC peaks were called using MACS2 v2.2.9.1. Compartments were called using cooltools v0.6.0. For ATAC analysis, reads over consensus peaks were counted from each sample using the featureCounts package v2.0.3. The resulting counts matrix was input into DESeq2 v1.38.3 for normalization and differential calls. Metagene plots were generated using the deepTools packages computeMatrix and plotProfile. Annotations ATAC peaks and loop anchors were generated using the ChIPSeeker R package. Differential compartment score analysis was performed using the limma package in R. PC1 eigenvectors were compared between KO and WT conditions using the empirical Bayes variance moderation. Significance was calculated using t-statistics and P-values were adjusted for multiple testing using Benjamini–Hochberg false discovery rate (FDR). Bins with adjusted P-values <0.05 were considered significant. Statistics for gene expression over compartment deciles were performed using a Wilcoxon rank-sum test followed by FDR correction.

#### RNA-Seq

RNA-Seq reads were processed using the Nextlfow nf-core/rnaseq v3.20.0 pipeline using default parameters. Reads were quality controlled using FastQC v0.11.9. Reads were aligned to the mouse GRCm38 genome using STAR v2.7.11b. Gene transcript counts were quantified using Salmon v 1.10.3. Normalization and differential analysis were performed using DESeq2 v1.38.3. Repeat expression was quantified using TEtranscripts v2.2.3 and repeat information was obtained from the UCSC genome browser’s Repeat Masker database.

## Data availability

Data published in this study is available through GEO under the following accession numbers: ChIP-seq GSE333216, HiCAR GSE333217, RNA-seq GSE333219.

## RESULTS

### Histone *H1f4* mRNA expression is reduced in postmitotic cells but maintained in differentiated neurons throughout the mouse brain

Many histone mRNAs lack polyA tails, thus they are not reliably represented in oligo dT primed sequencing datasets (21). To determine when and where *H1f4* is expressed in the mouse brain, we performed reverse transcription and quantitative PCR (qRT-PCR) for *H1* family genes on random hexamer-primed cDNA from mouse cortex, hippocampus, and cerebellum across brain development. Expression of all tested *H1* mRNAs was highest in embryonic development and decreased across all three brain regions with age (**Fig. 1**). Consistent with the expression levels of the *H1* mRNAs being highest in dividing cells, we found that *H1f1-H1f5* levels were already falling by birth in cortex and hippocampus (**Fig. 1B, C**). By contrast neurogenesis continues through the first week of postnatal life in cerebellum, and in this brain region H1 levels stayed high through P7 (**Fig. 1A**). Importantly, despite developmental downregulation, we continued to observe measurable expression of *H1* mRNAs throughout adulthood, with *H1f4* and *H1f0* mRNAs showing highest expression in all brain regions (**Fig. 1 D-F**).

**Fig. 1:**
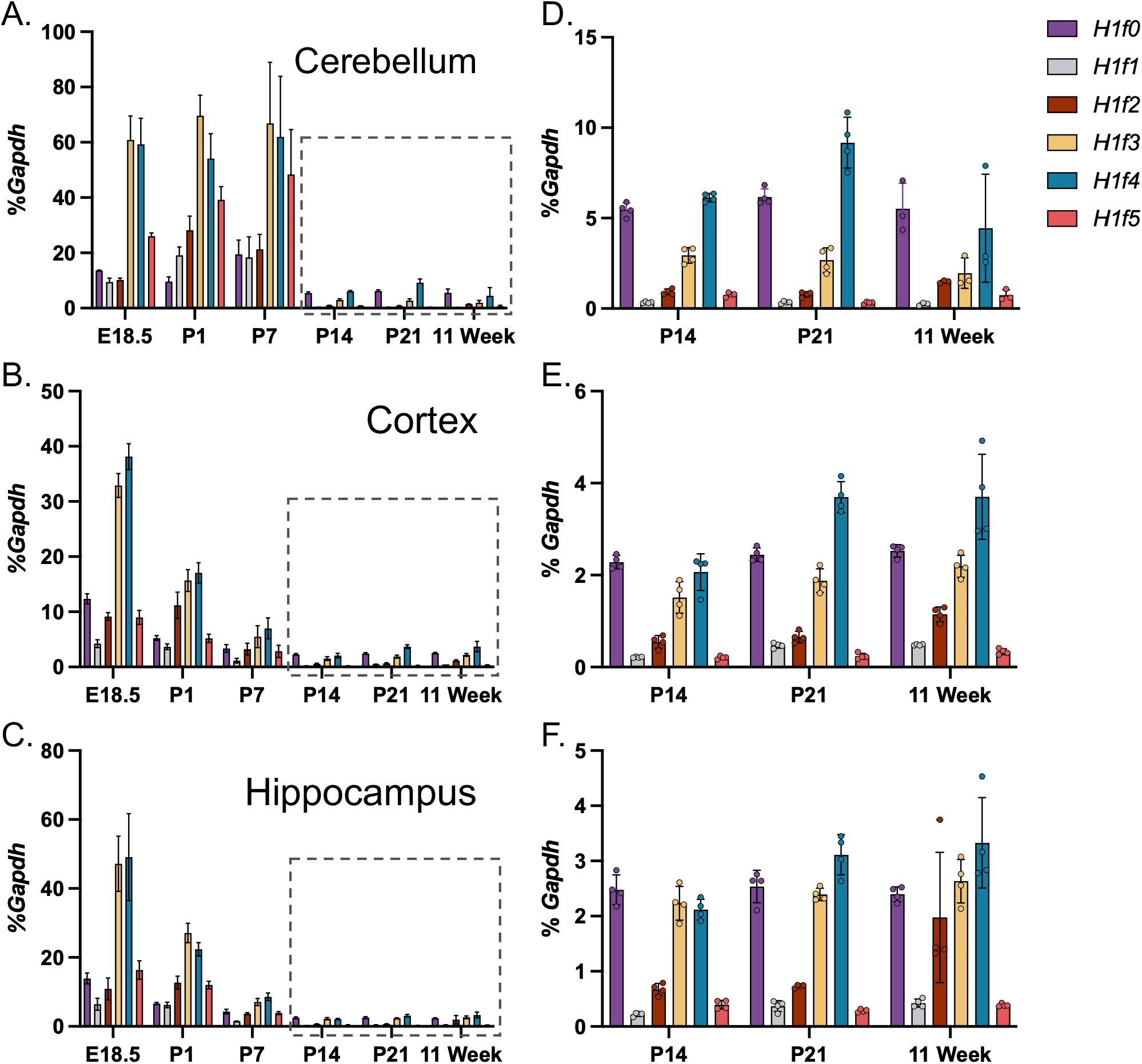
Somatic linker histone mRNAs are reduced but maintained in mature neurons. A-C: Expression of somatic linker histone mRNA in lysates from mouse cerebellum (A), cortex (B), and hippocampus (C) across developmental timepoints as measured by qRT-PCR. cDNA was made using random hexamer priming. D-F: Pullouts of A-C at postnatal ages after cessation of neurogenesis. Bars show mean and error bars are standard deviation.

By comparing cDNA primed with random hexamers to that primed with oligo-dT, we confirmed that the majority of linker histone mRNAs are detected only with random hexamer priming (**Fig. 2A,B**). By contrast, *H1f0* is equally detected by both methods of priming, and about 44% of *H1f2* transcripts appear in the oligo dT primed sample (**Fig. 2A,B**), suggesting polyadenylation of a subset of these transcripts, as previously reported (21). Linker histone *H1f1-5* mRNAs that are not polyadenylated end in a 3’ stem-loop recognized by the stem-loop binding protein (SLBP), which is required for their processing and translation (22). *Slbp* expression is highest during S-phase in dividing cells, but it has not previously been studied in the mouse brain. By qRT-PCR we confirmed that, although *Slbp* is highest during embryonic development at times of neurogenesis, it remains expressed in the adult mouse brain in cerebellum, cortex, and hippocampus (**Fig. 2C**).

**Fig. 2:**
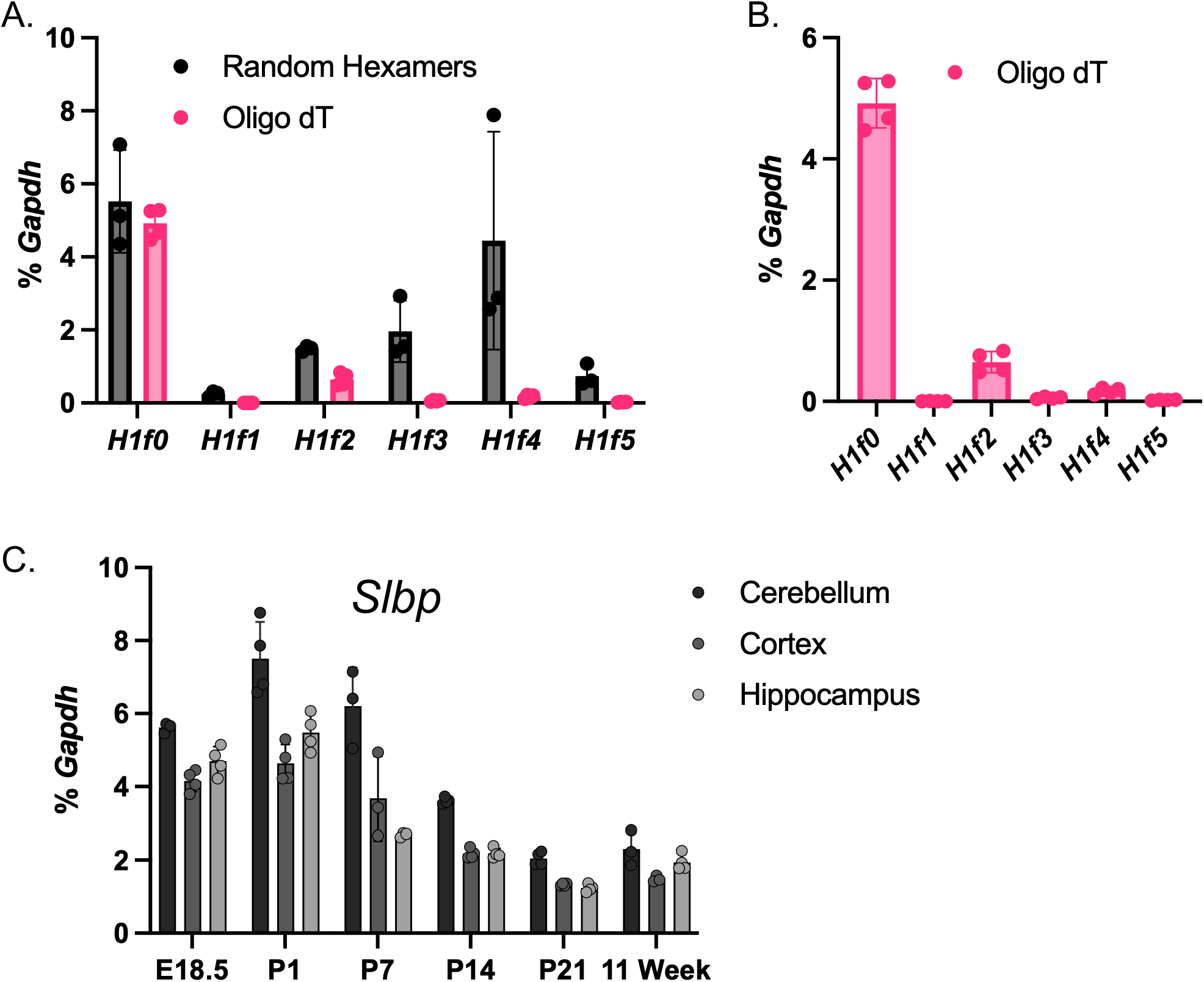
*H1f1-H1f5* are not polyadenylated in adult mouse brain. A: qRT-PCR for somatic linker histones from adult mouse cerebellum in cDNA made with either random hexamers (black) or oligo dT (pink) as indicated. B: Data from (A) regraphed with only oligo dT data shown for *H1f1-H1f5* to visualize relative expression. C: qRT-PCR for *Slbp* in the indicated brain regions and time points. Bars show mean and error bars are standard deviation.

Our qRT-PCR was conducted on bulk brain tissue, which contains many cell types. To confirm that *H1f4* mRNA is expressed in neurons, we performed RNA fluorescence in situ hybridization (FISH) for *H1* mRNAs on brain sections from mice (**Fig 3, Fig. S1**). We co-labeled cerebellar slices with RNAscope FISH probes for *H1f2* or *H1f4* together with a probe for *Map2*, a neuronal marker. These data confirmed that *H1f2* and *H1f4* signal is found in neurons at postnatal stages of development and into adulthood (**Fig. 3A**). We validated the specificity of the *H1f2 and H1f4* probes on cerebella from *H1f2/H1f4* double knockout (dKO) mice, confirming that we lost *H1f2* and *H1f4* signal, while preserving signal for *H1f3* and *Map2* (**Fig. 3B**). *H1f2* and *H1f4* transcripts also co-localized with *Map2* in cortex, and hippocampus taken from mice at P7, P21, and P60, confirming that these neurons continue to synthesize *H1* histone mRNA well after terminal differentiation (**Fig. S1**).

**Fig. 3:**
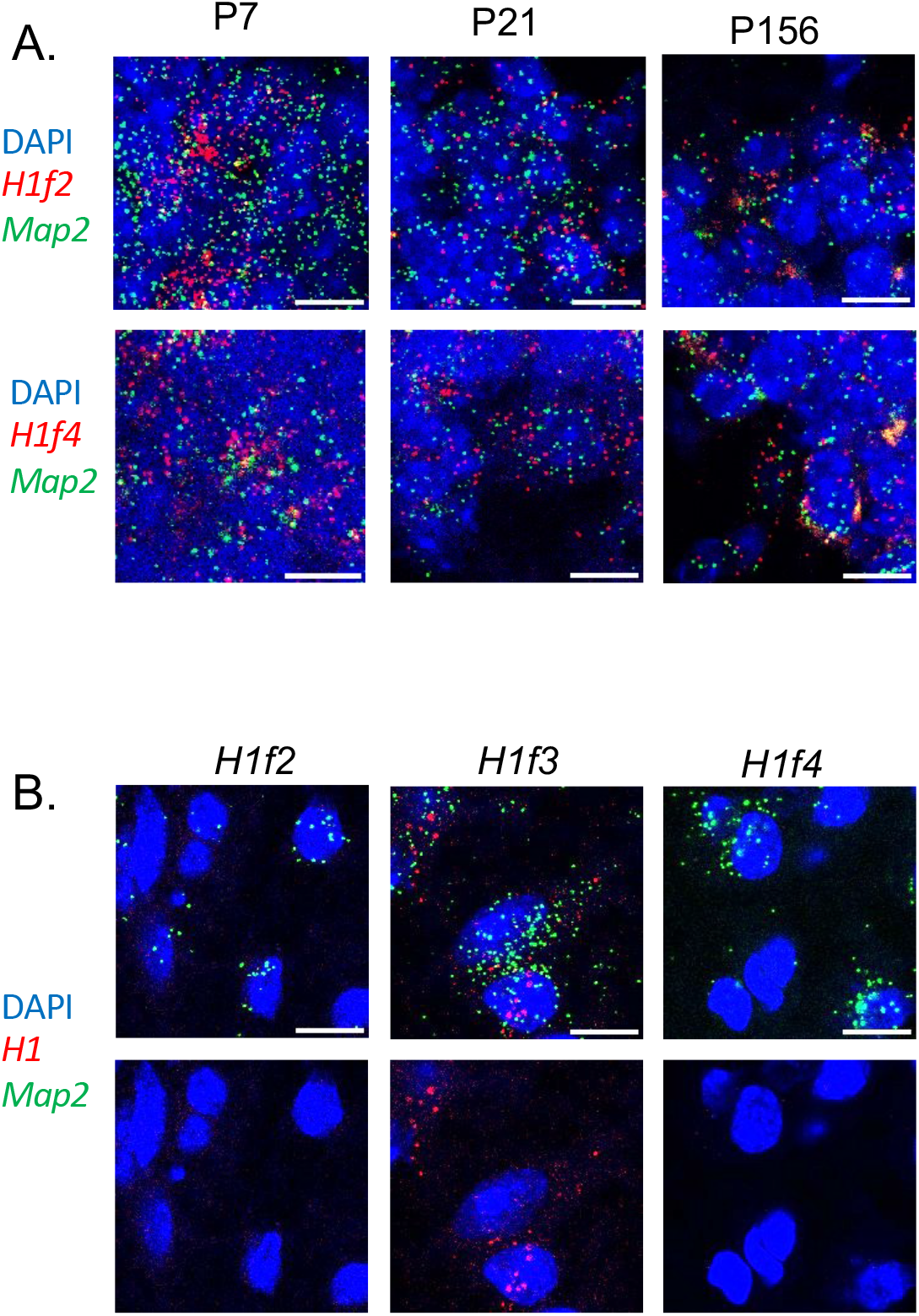
Histone H1 mRNA is found in postmitotic neurons of the mouse cerebellum. A: RNAScope for *H1f2* and *H1f4* in developing and adult cerebellum. DAPI marks nuclei and *Map2* is shown as a neuronal marker. B: Knockout validation of RNAScope fluorescence in situ probes for *H1f2* and *H1f4* using brain slices from *H1f2/H1f4* double knockout mice. *H1f3* is shown for comparison. Scale bar = 10µm.

### H1.4 protein is widely expressed in neurons and other cell types of the adult mouse brain

To determine which brain regions and cell types express histone H1.4 protein in the mouse brain, we generated an N-terminally tagged 3xFLAG-Myc-*H1f4* knockin mouse line, following a strategy that was used in previous studies to tag H1.0, H1.2, and H1.3 (23,24)(**Fig. 4A**, **S2**). This allowed us to use well validated epitope tag antibodies against FLAG and Myc to identify H1.4. To quantify the amount of H1.4 protein present in the mouse brain over time, we harvested tissue from cerebellum, cortex, and hippocampus of 3xFLAG-Myc-*H1f4* knockin mice or their WT littermates at postnatal day 7 (P7), P21, and P60. We acid extracted histones for western and blotted for H1.4, H1.0 and histone H3 as a loading control (**Fig 4B,C**). As reported previously, (5), we confirmed that H1.0 is minimally expressed at P7 and rapidly accumulates with brain maturation (**Fig. 4B,C**). H1.4 protein was robustly expressed at all timepoints tested, and in cortex and hippocampus we observed an increase in protein over time (**Fig. 4B,C**). This is notably in contrast to the decline in *H1f4* mRNA over this same time period. This suggests that post-transcriptional control of mRNA translation or changes in the rate of H1 protein turnover in the mature brain are crucial for controlling H1.4 protein levels in postmitotic neuronal chromatin.

**Fig. 4:**
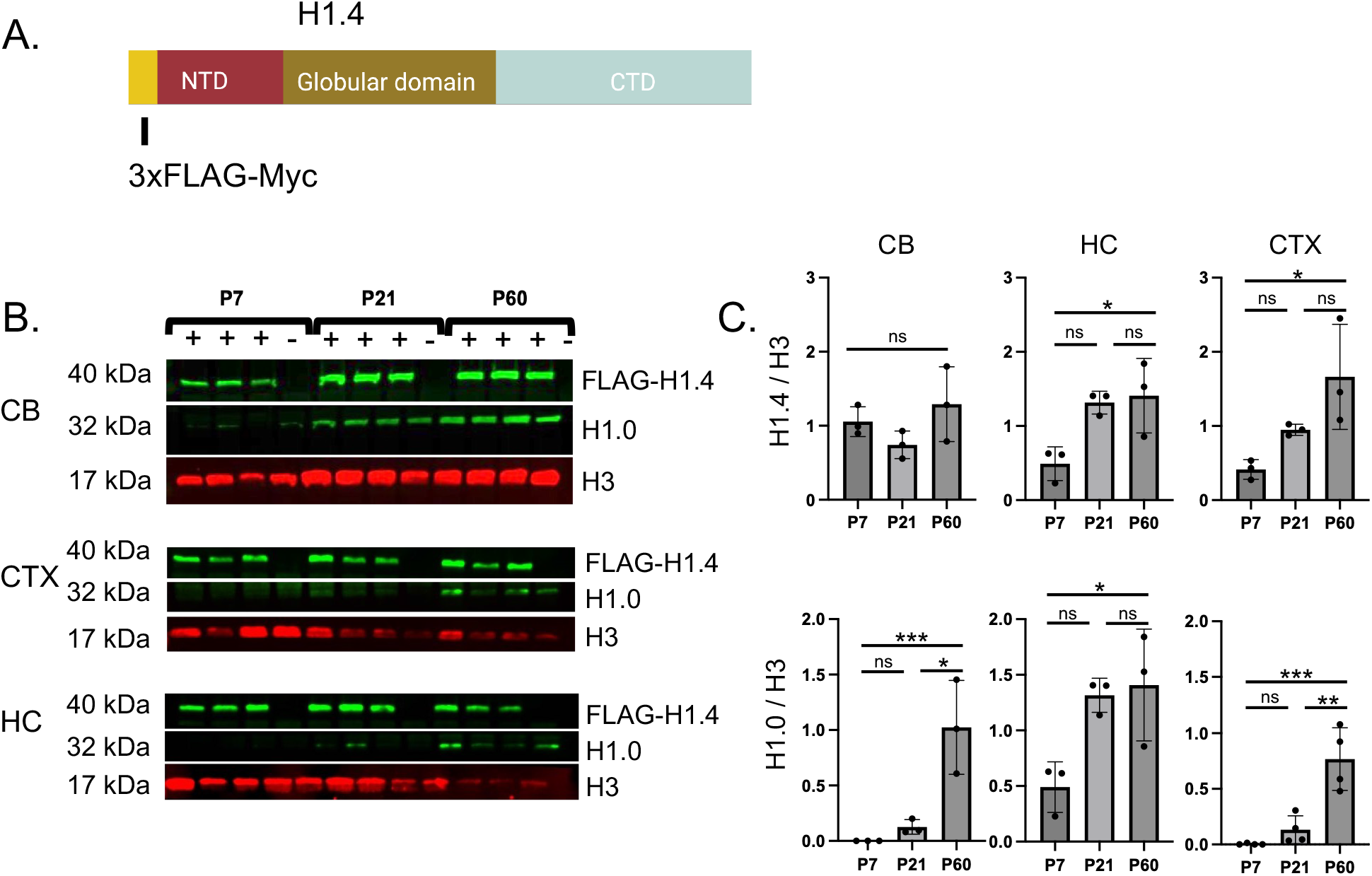
Histone H1.4 protein accumulates in the mature mouse brain. A: Diagram of the 3xFLAG-Myc-H1.4 protein made from the *H1f4* Tag knockin line. B: Western blot of acid extracted histones from heterozygous 3xFLAG-Myc-*H1f4* mice or their WT littermates. 3xFLAG-Myc-*H1f4* genotype is shown at top (+ epitope tagged, - WT). Antibodies were against the FLAG epitope for H1.4. Antibodies against H1.0 were used for comparison and histone H3 as a loading control. Tissues from cerebellum (CB), cortex (CTX), and hippocampus (HC). C: Quantification of western blots for 3xFLAG-Myc-H1.4 (top) and H1.0 (bottom) normalized to histone H3 in each lane. n=3 biological replicates of the 3xFLAG-Myc-*H1f4* mice and 1 WT mouse per timepoint. Bars show mean and error bars are standard deviation. * p<0.05, **p<0.01, ***p<0.001 one-way ANOVA for time.

To visualize H1.4 expression in different developmental stages and cell types of the mouse brain, we performed immunofluorescence on brain sections taken from 3xFLAG-Myc-*H1f4* KI mice. At low power, we observed that H1.4 is broadly and mostly evenly expressed throughout the adult brain (**Fig 5A**). In the cerebellum, at P7 the external granule layer (EGL) is comprised of both cerebellar granule neuron (CGN) progenitors and newborn neurons. Newborn CGNs then migrate to the internal granule cell layer which, at both P7 and P60, is composed almost entirely of Zic1/2-expressing postmitotic, mature CGNs. We saw expression of H1.4 in CGNs at both ages with strongest H1.4 staining in the IGL, suggesting that H1.4 protein accumulates in these neurons after they exit the cell cycle and reach their final stages of maturation (**Fig 5B**). Cerebellar expression of H1.4 is not limited to CGNs, because calbindin-expressing Purkinje neurons also stained positive for H1.4 at both P7 and P60 (**Fig 5C**).

**Fig. 5:**
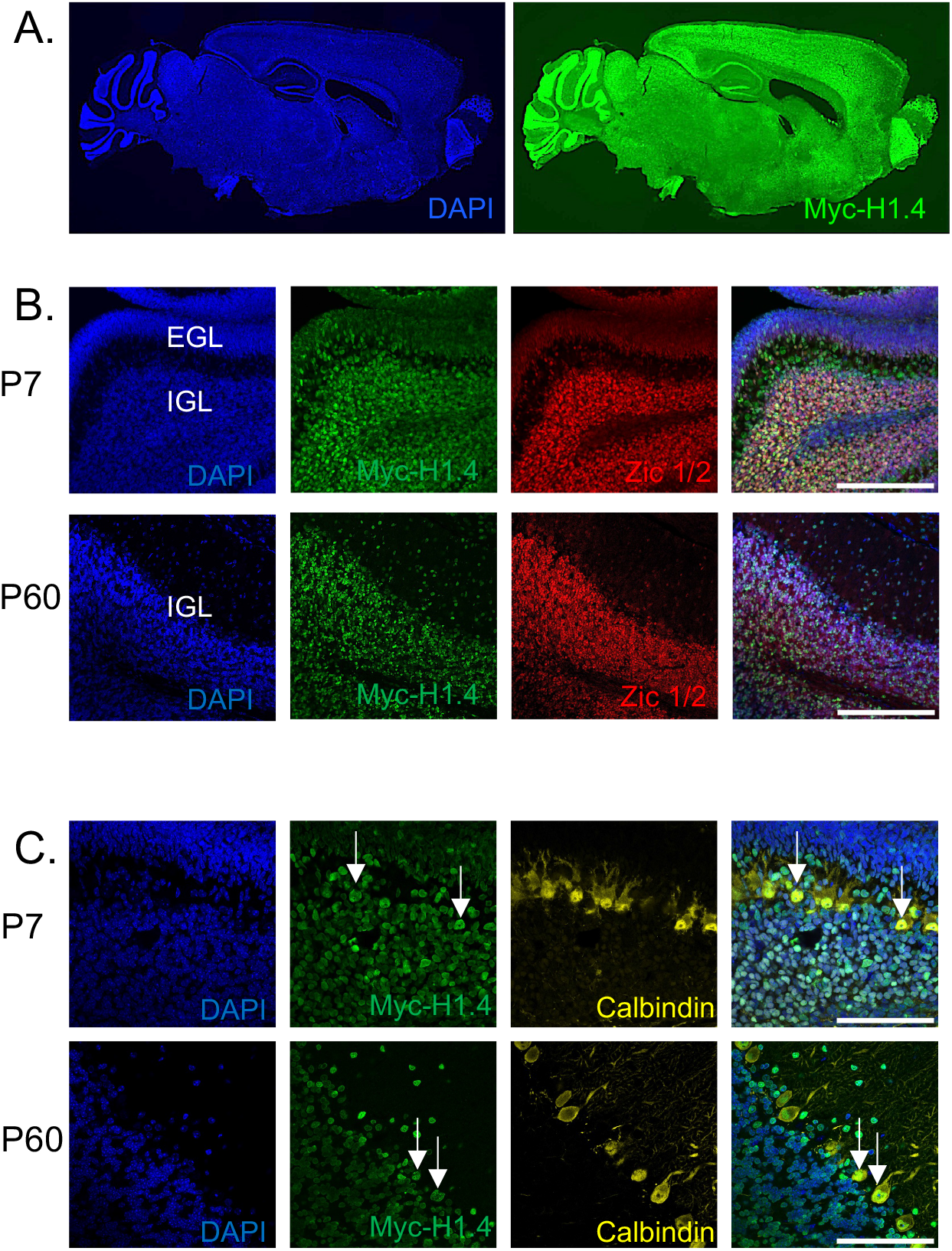
Histone H1.4 accumulates in mature neurons across the brain. A: Immunofluorescence images of parasagittal sections from P60 3xFLAG-Myc*-H1f4* knockin mouse brains. Antibodies against the Myc tag were used for H1.4. DAPI labels nuclei for anatomical orientation. B: Immunofluorescence images of P7 (top) and P60 (bottom) parasagittal cerebella sections. Zic1/2 is a marker for cerebellar granule neurons (CGNs). P7 shows both the external (CGN progenitor) and internal (CGN) granular layers. P60 shows the internal granular layer. Scale bar = 200µm. (C) Co-immunofluorescence images of H1.4 with calbindin positive cells in the cerebellum of P7 and P60 mice. Calbindin is a marker for Purkinje neurons. Nuclei of representative Purkinje neurons indicated with arrows. Scale bar = 100µm.

In the adult cortex and hippocampus, H1.4 is widely expressed, overlapping but also extending beyond the distribution of the methyl-DNA binding protein MeCP2, a neuron-enriched chromatin regulatory factor that has potential functional and biochemical interactions with histone H1 (23,25–27) (**Fig. 6A**). In hippocampus, H1.4 colocalized with MeCP2 in dentate gyrus and neurons of area CA1 (**Fig. 6B, C**). H1.4 was also present in some cells with low MeCP2 expression and elongated nuclei that are likely to be astrocytes (**Fig. 6C**). However in the same field, H1.4 negative, MeCP2 negative nuclei are also visible suggesting that H1.4 expression is not universal in all brain cell types. In the cortex, we found that IBA1-expressing microglia (**Fig. 6D**) and GFAP-positive astrocytes (**Fig. 6E**) express H1.4, although microglia in particular appeared to express much lower levels of H1.4 compared with other cells we labeled (**Fig. 6D**). Finally to determine the subnuclear distribution of H1.4, we took high magnification confocal images through hippocampal CA1 neurons co-stained for H1.4 and MeCP2 (**Fig. 7**). MeCP2 is highly enriched at chromocenters, which are dense sites of heterochromatin comprised of condensed pericentromeric satellite DNA. H1.4 shows some overlap with these bright spots of DAPI and MeCP2 signal, however overall it has a broader distribution throughout the nucleus with some enrichment at the nuclear envelope. These images are similar to the reported distribution of H1.4 in human breast cancer cells (28), supporting that the epitope tag does not alter the endogenous protein distribution.

**Fig. 6:**
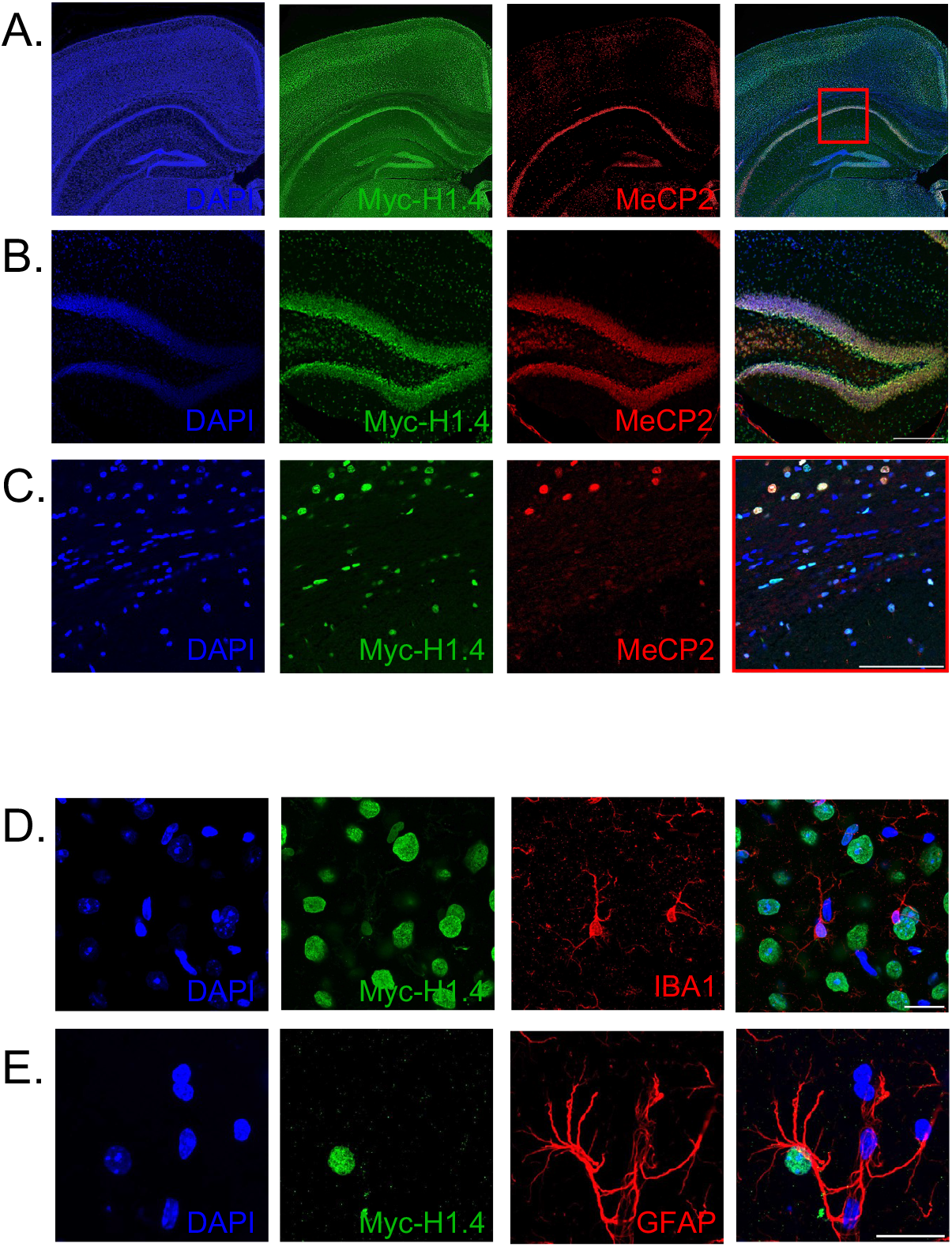
Histone H1.4 is expressed in both neurons and non-neuronal cells in the adult mouse cortex and hippocampus. A) Immunostaining of epitope tagged histone H1.4 with an anti-Myc antibody on a coronal section through the hippocampus and cortex. MeCP2 is shown for comparison as a broadly expressed, neuron-enriched, chromatin binding protein. B) shows the dentate gyrus, scale bar = 200µm. C) shows CA1, scale bar = 100µm. Co-labeling of tagged H1.4 (Myc) with a marker for D) microglia (IBA1) or E) astrocytes (GFAP). Scale bar = 20µm. DAPI labels nuclei for anatomical orientation.

**Fig. 7:**
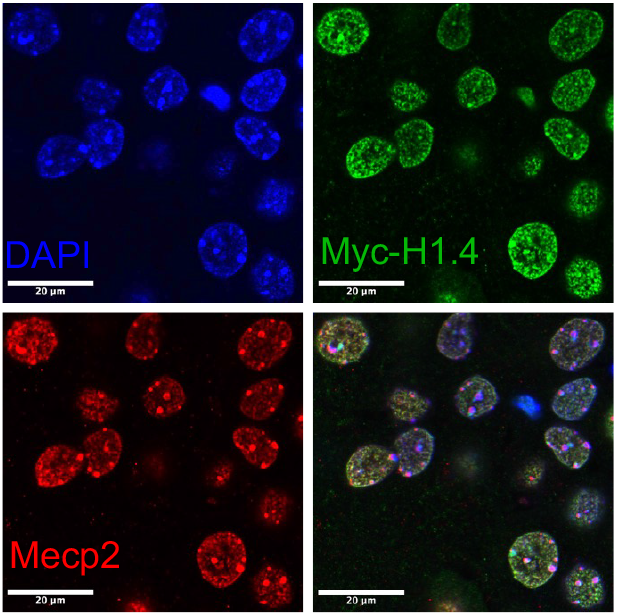
H1.4 has a subnuclear distribution in CA1 hippocampal neurons that is distinct from MeCP2. 40X confocal images of tagged H1.4 in CA1 hippocampal neurons of the adult mouse brain. Antigen retrieval was used to maximize nuclear signal. MeCP2 is shown for comparison as a nuclear protein that is highly concentrated at chromocenters visible with DAPI. Scale bar 20µm.

### The genomic distribution of H1.4 shifts toward repressed chromatin as neurons mature

The H1 histones in general have been thought to bind broadly across the genome with greater enrichment in heterochromatin (4,29,30). Only histone H1.0 has previously been profiled for its genomic distribution in neurons, and it showed a broad heterochromatic distribution similar to the methyl-DNA binding repressor MeCP2 (23). To determine the genomic distribution of H1.4 in neurons, we performed chromatin immunoprecipitation followed by sequencing (ChIP-seq) for histone H1.4 from our 3xFLAG-Myc-*H1f4* knockin mice using an anti-Myc antibody. Importantly we confirmed that there was no significant chromatin pulldown using the anti-Myc antibody from cerebellum of WT mice (**Fig. S4A)**. We compared H1.4 binding in granule neuron progenitors purified from P7 cerebellum to ChIP-seq from cerebellum of adult (P60) mice, which is almost entirely composed of mature CGNs (**Fig. 8**). Input-normalized ChIP peaks were highly replicable between samples at each age; however, we observed a dramatic shift in H1.4 distribution between GNPs and the mature cerebellum (**Fig. 8A**). In GNPs, the H1.4 distribution included active regions of the genome marked by histone H3K27ac, whereas in the mature cerebellum H1.4 much more closely tracked the heterochromatic distribution of MeCP2 reported from forebrain neurons (**Fig. 8B**). This temporal distinction in H1.4 distribution within active regions of the genome can also be seen locally at transcription start sites (TSS), as revealed by the relative enrichment of H1.4 at active TSSs in GNPs and the relative de-enrichment of H1.4 at active TSSs in adult cerebellum (**Fig. 8C,D**).

**Fig. 8:**
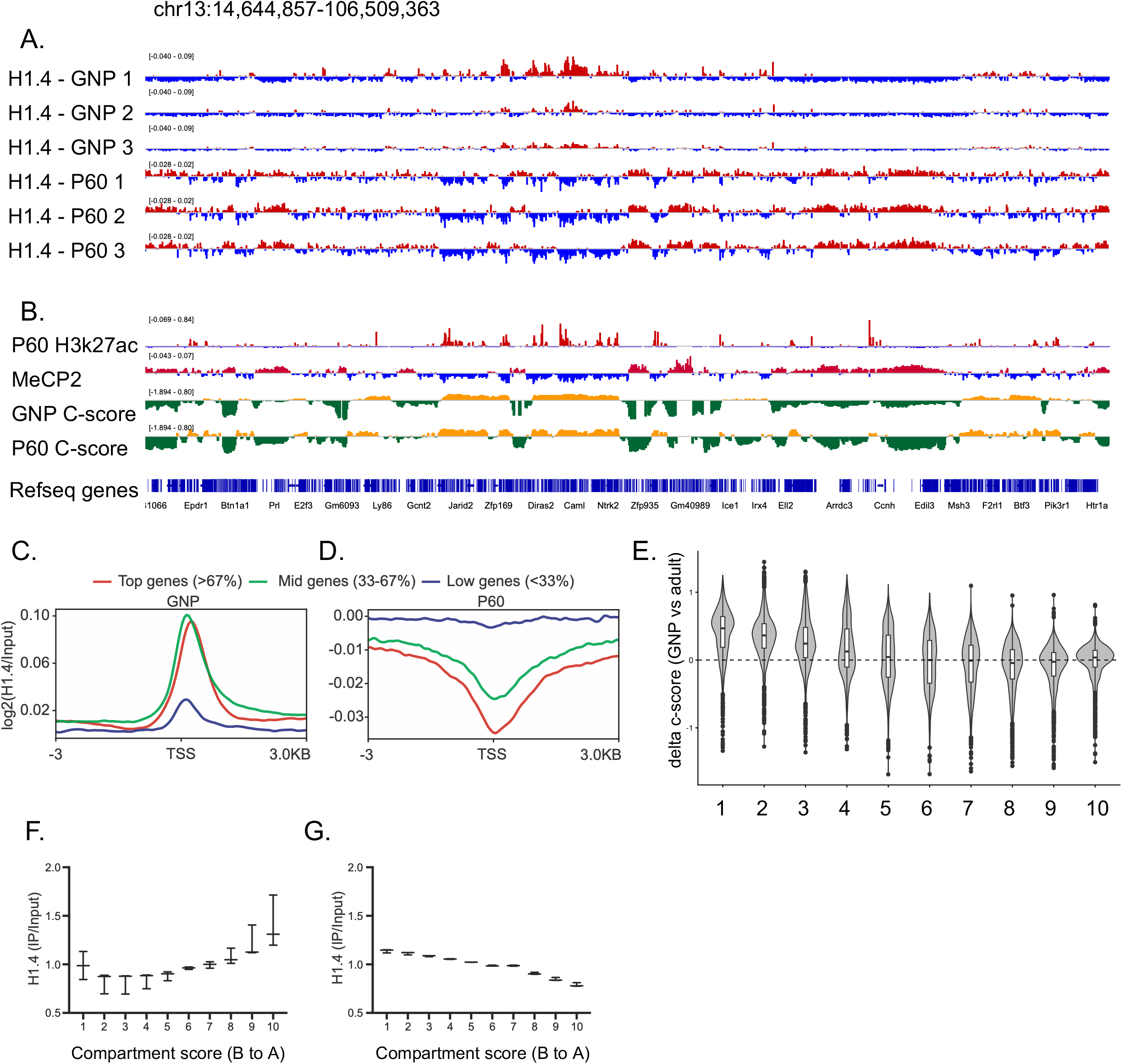
H1.4 accumulates in heterochromatin when CGNs mature. A: Genome viewer track of 3xFLAG-Myc-H1.4 ChIP-seq data from purified P7 granule neuron progenitors (GNPs) or adult cerebellum (P60) aligned to mouse chromosome 13. All ChIP signals are shown subtracted from their respective input samples. Regions of enrichment in red; regions of de-enrichment in blue. B: H3K27ac ChIP-seq data are from adult cerebellum (31) and MeCP2 ChIP-seq data are from adult mouse forebrain excitatory neurons (23). Compartment scores derived from HiCAR chromatin conformation data in GNPs or P60 cerebellum. The Eigenvalues are represented in orange for A compartment/euchromatin, and green for B compartment/heterochromatin. C,D: Myc-H1.4 ChIP read distribution +/-3kB from the transcription start site (TSS) of genes binned by relative expression level in GNPs or P60 cerebellum. E: Shift in compartment score between GNP and P60. 100kb genomic bins were ranked by P60 compartment score and separated into deciles (1-10, B to A). Boxes represent median and interquartile range, whiskers represent 1.5xIQR, with outliers plotted as individual points. F,G: Distribution of H1.4 ChIP reads from GNPs or P60 cerebellum. 100kb bins were ranked by GNP or P60 compartment score and separated into deciles (1-10, A to B). ChIP reads were counted within each decile and normalized against respective inputs, n=3 per group.

### Developmental chromatin compaction is impaired in CGNs from *H1f2/H1f4* double knockout mice

Our ChIP-seq data show that H1.4 is distributed over large megabase scale regions of the genome, suggesting its distribution may track with chromatin compartments. To identify chromatin compartments in both GNPs and mature CGNs, we used a chromatin conformation method called HiCAR (HiC on Accessible Regulatory DNA). HiCAR uses Tn5 transposase to identify chromatin interactions anchored at sites of accessible chromatin, providing information on both chromatin accessibility and chromatin architecture(19). The genomic distributions of the HiCAR 1D open chromatin peaks were highly similar between samples and shared the genomic distribution expected for identification of both promoters and enhancers **(Fig. S4B)**.We confirmed that the R2 HiCAR reads, which arise from Tn5 transposition, overlapped sites we previously called as DNAseI hypersensitive open chromatin in GNPs and CGNs (31), whereas the proximity ligated R1 reads lacked this enrichment **(Fig. S4C)**. Because HiCAR loops are anchored at one end on open chromatin, they are enriched for transcriptional regulatory interactions relative to other HiC methods (19), and we confirmed that loops showed the expected promoter-centric distribution **(Fig. S4D)**. Importantly, consistent with a study that used a different chromatin conformation method to follow CGN chromatin across development and maturation (32), we found that DNA within heterochromatic regions of adult CGNs was significantly more compact than the same regions of the genome in GNPs **(Fig. 8E**). Thus these data show that we can use HiCAR to quantitatively evaluate the developmental regulation of chromatin compartments in CGNs.

When we aligned the H1.4 ChIP-seq tracks against the HiCAR compartments, these data revealed that H1.4 has significant overlap with A-like regions in GNPs, whereas binding more closely tracks the heterochromatic B domains in adult cerebellum (**Fig. 8A,B**). When we graphed H1.4 distribution versus genome bins ranging from most B-like to most A-like, the relative enrichment of H1.4 in A domains in GNPs versus B domains in CGNs could be seen at a global scale (**Fig. 8F,G**).

To address whether H1.4 is important for maintenance of chromatin compartments, we performed HiCAR on cerebella from adult *H1f2/H1f4* dKO mice compared to adult WT (**Fig. 9A**). HiCAR libraries from the dKO mice had similar open chromatin and loop anchor distributions compared with the GNP and WT adult cerebellar libraries (**Fig. S4B**). We saw minimal differences in HiCAR 1D open chromatin between WT and dKO samples and very few significant changes in 3D chromatin looping (**Fig. S5A,B**). 1D HiCAR open chromatin peaks and regions of active gene expression aligned well with A compartment designations in both genotypes (**Fig. S5C**). However we did find multiple regions across the genome where chromatin compartments were significantly different between WT and dKO mice (**Fig. 9A,B**). Specifically we identified 1005 compartment bins that became significantly less compact in the dKO mice compared with WT and 129 that became more compact (**Fig. 9C**). When we graphed the change in chromatin compaction in bins ranging from most B-like to most A-like chromatin from WT mice, we observed that decompaction in the dKO mice was most extensive in the strongest B-compartment regions (**Fig. 9D**). Indeed, most of the significant compartment shifts involved changes in the magnitude of B compartment chromatin compaction, whereas only a few regions switched from B to A or A to B (**Fig. 9E**). Interestingly, the regions that fail to compact in dKO mice relative to WT are also regions that tend to be more compact in mature CGNs relative to GNPs (**Fig. 9F,G**). Thus these data suggest that loss of H1.2 and H1.4 leads to a failure of developmental maturation of chromatin compaction in CGNs.

**Fig. 9:**
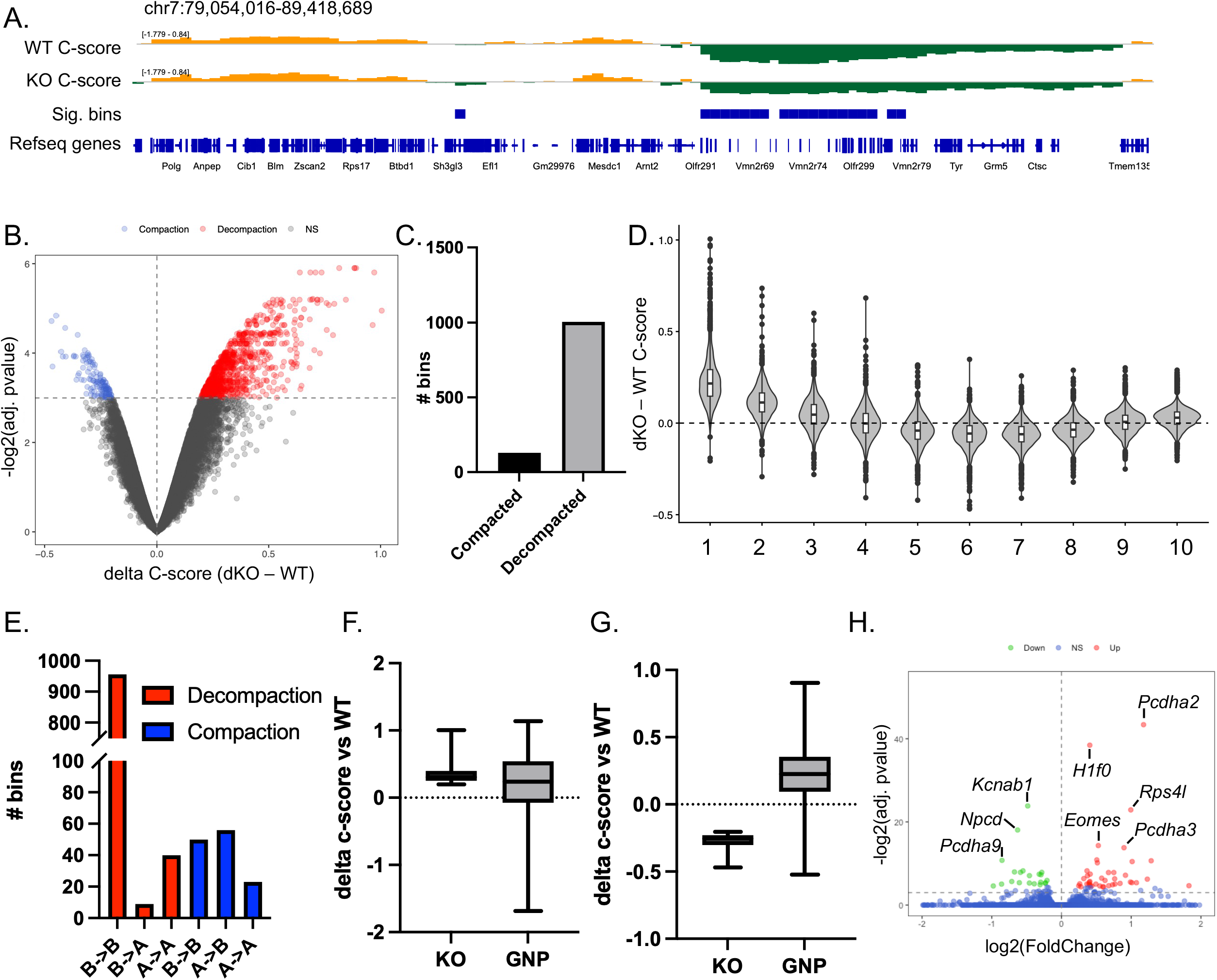
Double *H1f2/H1f4* knockout mice show impaired condensation of B compartment chromatin. A) Representative example of chromatin regions showing significant changes in chromatin compartment score in adult cerebellum of WT vs *H1f2/H1f4* double knockout (KO) mice. The Eigenvalues are represented in orange for A compartment/euchromatin, and green for B compartment/heterochromatin. B) DEseq quantification of chromatin regions that are significantly more A (red) versus more B (blue). C) Number of compartments that are more compacted or decompacted in cerebellum of KO versus WT mice. D) Shift in compartment score between KO and WT. 100kb genomic bins were ranked by WT compartment score and separated into deciles (1-10, B to A). Boxes represent median and interquartile range, whiskers represent 1.5xIQR, with outliers plotted as individual points. E) Breakdown of compartment transitions for genome regions with significantly altered chromatin compaction. F,G) Comparison of 100kb genomic bins that either gain or lose compaction in KO vs WT and GNP vs WT. H) Volcano plot of DESeq quantification of differential gene expression in KO vs WT mouse cerebellum.

Finally, to determine whether the loss of H1.2 and H1.4 affects gene expression, we performed bulk RNA seq on adult cerebellar cortex from *H1f2/H1f4* dKO and WT mice. We observed relatively few changes in gene expression, with 24 genes showing reduced expression and 44 genes showing gained expression in dKO cerebellum compared with WT (**Fig. 9H**). Among the upregulated genes, we found several histones including the linker histone *H1f0*, which is consistent with prior evidence that members of the H1 histone variant family compensate for the loss of other family members (8,14,33). To determine how changes in gene expression related to the compaction changes in dKO mice, we quantified gene expression in WT and dKO mice within compartment bins from most to least compact (**Fig. S5D**). These data show the general correlation of higher gene expression with A compartments (**Fig. S5C,D**), however there were no significant differences in levels of overall gene expression between WT and dKO mice in any of the bins (**Fig. S5E**). When we mapped differentially expressed genes into the compartment bins, we found they were mostly in the weak A compartments, whereas compaction differences were in strong B. These data raise the question of how changes in chromatin compartments may influence gene expression, which we discuss below.

## DISCUSSION

In this study, we determined the developmental, regional, and genomic distribution of the Rahman Syndrome gene and linker histone H1.4 in the mouse brain. We confirmed that *H1f4* is expressed in postmitotic neurons as well as dividing neural progenitors, and that H1.4 protein continues to accumulate in the adult brain despite falling levels of mRNA expression. Using ChIP-seq, we observed that H1.4 associates with active genes in neural progenitors but accumulates in heterochromatic B compartment chromatin in mature neurons. Consistent with a role for H1.4 in chromatin condensation, our chromatin conformation assays revealed that B compartment chromatin fails to reach normal levels of compaction in the cerebellum of adult *H1f2/4* dKO mice. These data offer a comprehensive characterization of Histone H1.4 in the brain and build a foundation for future studies that address how H1.4 disruption leads to neurodevelopmental phenotypes in Rahman Syndrome.

### H1.4 protein accumulates in non-dividing cells of the adult brain including neurons

H1.4 is termed one of the “replication dependent” histones both because its RNA is most highly expressed during S-phase and because the processing and translation of this mRNA depends on the S-phase enriched protein SLBP (22,34). In the brain, levels of the replacement histone H1.0 rise sharply during the first weeks of postnatal development, suggesting that the polyadenylated transcript encoding this histone is an important source of newly synthesized H1 in maturing neurons (5). However even early studies of histone H1 expression in dividing and nondividing cells of the mouse liver, kidney, lung, and thymus indicated that H1.4 accumulated in non-diving somatic cells (2), and in nuclear extracts from rat cerebral cortex, H1.4 was observed to increase in abundance relative to other H1s across brain development (5). To track histone H1.4 expression both quantitatively and at the single cell level, we developed an N-terminal 3xFLAG-Myc tagged *H1f4* knockin mouse. Our data reveal that histone H1.4 is found in most, though perhaps not all, cells of the developing and adult mouse brain, and that it is expressed in postmitotic neurons well into adulthood, long after cell division has ceased.

Histones are among the most long-lived proteins in the brain. Whereas the median lifetime of brain proteins was estimated by isotopic pulse-chase labeling to be about 10.7 days, the lifetime of H1.4 was calculated at 30 days in adult mouse cortex and 97 days in cerebellum (35). Among the H1 family members, H1.0 and H1.3 also have long lifetimes, whereas the turnover of H1.2 is much faster, with a calculated lifetime of 14-26 days in different brain regions. Although these lifetimes are long, the accumulation we observed of histone H1.4 protein in adult mouse neurons implies that there remains some ongoing synthesis and incorporation of H1.4 into postmitotic chromatin. Consistent with this prediction, our FISH data show that we continue to see *H1f4* mRNA in neurons, as well as *H1f2* and *H1f3*, even at the latest time points examined.

We do see varying levels of H1.4 expression in different brain cell types. In particular microglia showed weaker immunostaining for histone H1.4 compared with neurons despite having highly compact nuclei that are similar in size to CGNs. Given that mRNA is not a good indicator of H1 protein levels, future studies that isolate specific brain cell types for quantitative mass spectrometry or western blotting analysis will be most useful for determining the H1 family composition of different brain cell types.

### Histone H1.4 accumulates in repressed B-compartment chromatin in the adult brain

In addition to showing where H1.4 is expressed in the brain, we used the epitope tag on our knockin allele to determine the distribution of H1.4 across the genome. As expected for a histone, our ChIP-seq data show that H1.4 has a broad genomic distribution, with a pattern of enrichment and de-enrichment that tracks the megabase scale of chromatin A/B compartments. Strikingly, we found that the pattern of H1.4 binding was significantly different when we compared P7 granule neuron progenitors, which are still dividing, with postmitotic cerebellar granule neurons from P60 brains. In adult cerebellum, histone H1.4 was enriched in the most compact B-like chromatin regions, mirroring the heterochromatin distribution of histone H1.0 and MeCP2 (23). By contrast in GNPs, histone H1.4 was distributed across a broad range of chromatin compartments including in active chromatin domains enriched for H3K27ac, and it showed binding at transcription start sites of expressed genes.

Only one prior study has reported ChIP-seq for endogenous histone H1.4 as part of a systematic comparison of six H1 family members in the human breast cancer cell line T47D-MTLV (30). The authors reported that whereas H1.0, H1.2, H1.3, and H1.5 were enriched in the inactive B-compartments of chromatin, H1.4 and H1.10 were found in active A compartments. Our data are the first to compare the genomic binding pattern of H1.4 between differentiation states of the same cell type, and they suggest that the distribution of H1.4 changes when cells exit the cell cycle, raising the question of the mechanism that leads to this redistribution. Similar to other histones, the tails of the H1 histones can be post-translationally modified to alter their function and chromatin binding affinities (36–40). Top-down proteomics data show that histone H1.4 undergoes progressive phosphorylation at several Ser/Thr sites during mitosis (41). In one report, phosphorylation of H1.4 has been shown to promote its general dissociation from chromatin (42), though in others the phosphorylation of H1.4 is regulated locally at specific genes and associated with active transcription (43–45). In either case, we hypothesize that the regulation of H1.4 phosphorylation in differentiating CGNs could contribute to genomic redistribution of this histone in maturing neurons.

### Histone H1.2 and H1.4 are required for proper chromatin compaction in mature neurons

One of the most interesting questions about H1 histones is whether or not specific members of this gene family have distinct functions (46). Knockout of any single *H1* in mice shows minimal phenotypes, confounding understanding of their unique functions (7).Some of the challenges arise from intrafamilial compensation as loss of one *H1* gene leads to upregulation of others (7). Only triple knockout of *H1f2, H1f3,* and *H1f4,* which lowers overall H1 protein to 50% of normal, results in serious developmental defects (8).

Although the biochemical details of how specific H1s functionally differ remain to be fully understood, a growing body of data do suggest that the consequences of H1 regulation of chromatin differ by cell type. Detailed examination of knockouts in specific tissues has consistently supported a role for H1 proteins in the organization of A/B chromatin compartments in contexts that are relevant for specific cellular physiology. For example, (47) made triple conditional *H1f2, H1f3,* and *H1f4* knockouts in hematopoietic cells and showed that the resulting loss of H1 in CD8+ T cells led to selective decompaction of chromatin in two kinds of heterochromatin: both the most B-like compartments, which are constitutive heterochromatin marked by H3K9me3, as well facultative heterochromatin in the A compartment that is regulated by polycomb and marked by H3K27me3. These changes were associated with a decreased capacity for proliferation and a shift toward expression of T cell activation genes, showing the cell-intrinsic functional importance of chromatin compaction for T cell physiology. Similar chromatin architecture and histone modification landscape changes were seen in germinal center B cells lacking *H1f2* and *H1f4,* however in these cells the derepression of stem cell genes led to tumor formation, establishing these H1 variants as important tumor suppressors in this context (9). These data suggest that the particular complement of H1 histones expressed in a given cell type as well as the types of genes that are repressed by heterochromatin in that cellular context will determine the functional importance of H1s in cell physiology.

Neurons exit the cell cycle especially early in their lifespan, and perhaps as a result they regulate their chromatin in somewhat different ways from other cells. For example, neurons have extensive DNA methylation at CpA dinucleotides in addition to the more widely used methylation at CpGs (48) and some neurons centrally aggregate their heterochromatin rather than concentrating it at the nuclear periphery (49,50), Rod photoreceptors are one of the neuronal cell types with inverted heterochromatin that increases during postnatal development contemporaneously with increased histone H1.2 expression. This developmental regulation of rod heterochromatin is impaired in *H1f2/H1f4* double- and *H1f0/H1f2/H1f4* triple-knockout mice showing the general importance of histone H1s in heterochromatin formation regardless of intranuclear location (4). CGNs are similar to rod photoreceptors in having very compact nuclei, though they have the more common peripherally localized heterochromatin. Interestingly, CGNs continue to refine their chromatin architecture long after cell fate commitment is complete, showing a lifelong chromatin restructuring (32). Over time these cells show a shift in the magnitude of compaction such that heterochromatic regions become more compact and euchromatic regions become less compact. It is proposed that this strengthening of chromatin compartments over time permits the robust expression of developmentally late expressing genes that confer mature synaptic properties on CGNs in the adult brain (32). Our data show that developmental chromatin compaction in CGNs is impaired in *H1f2/H1f4* double knockout mice, raising the possibility that chromatin regulation by H1s may help to orchestrate functional maturation of these cells. It is important to note that the majority of altered chromatin architecture events we observed were decompaction in the most B-like heterochromatin, which likely explains why we saw minimal effects on gene expression. In addition, we observed upregulation of *H1f0* in our *H1f2/H1f4* double knockout mice, which may functionally compensate for the loss.

In summary, our data provide novel understanding of the expression, regulation, and function of Histone H1.4 in the brain. These data reveal that H1.4 has important functions in mature neurons, including the global organization of chromatin into active and inactive compartments, which is thought to be important for robust control of proper gene expression. Future studies that assess how chromatin changes in the presence of the Rahman Syndrome mutations in H1.4 may help to explain how changes in chromatin architecture can lead to intellectual disability in neurodevelopmental disorders.

## Acknowledgements

This work was supported by NIH grant R01NS136375 (A.E.W.) and a grant from the Simons Foundation Autism Research Initiative, SFARI (A.E.W.). We thank Gary Kucera and the Duke University Mouse Transgenic Core Facility for generation of the *H1f4* knockin mouse.

## Supplemental figures

**Fig. S1:** Somatic linker histone H1 mRNA in postmitotic neurons of cortex and hippocampus. RNAScope FISH for *H1f4* and *H1f2* in developing and adult cortex (A,B) and hippocampus (C,D). DAPI marks nuclei and *Map2* is shown as a neuronal marker. Scale bar = 10µm.

**Fig. S2:** Generation of 3xFLAG-Myc-*H1f4* knockin mouse strain. A: CRISPR knockin strategy for 3xFLAG-Myc tag. The single exon *H1f4* gene on chromosome 13 is shown. Tags were cloned in frame directly downstream of the translation start site of H1.4 in the donor vector. sgRNA cut sites were selected to be beyond the 3’ stem loop sequence. B: Inside to outside PCR of both the 5’ and 3’ ends of *H1f4* validated successful transgene knockin in ES cell line G2 (labeled in red). C: PCR using primers flanking the translation start site in n=2 heterozygous knockin mice and n =1 WT mouse.

**Fig. S3**. Uncropped western blots for 3xFLAG-Myc-*H1f4* mice. The full westerns from Figure 4 are shown uncropped here. We probed each single western with antibodies for each of the three targets.

**Fig. S4.** Validation of HiCAR chromatin conformation data from *H1f2/H1f4* dKO mice. A: ChIP qRT-PCR from WT and 3xFLAG-Myc mice. Bars show mean and error bars are standard deviation, n = 3 mice per genotype. B: Genomic annotation of 1D ATAC peaks with respect to genomic landmarks for GNP, WT P60, and KO P60. C: Sequence depth normalized (counts per million, CPM) of HiCAR R1 (ligated) and R2 (1D ATAC) reads plotted as signal coverage ±3kb around DNAse hypersensitive sites from (31). D: Genomic annotation of loop anchors for GNPs, P60 WT, and P60 KO.

**Fig. S5.** Comparison of gene expression and HiCAR data from *H1f2/H1f4* dKO mice A: MA plot of 1D ATAC peaks showing comparison of KO vs WT from DESeq. B: Volcano plot showing significant changes in chromatin interactions between KO and WT. C: Example genomic view of compartment scores, 1D ATAC peaks, and RNAseq data from HiCAR in WT and *H1f2/H1f4* dKO mice. D: Boxplot of gene expression relative to compartment score. 100kb genomic bins were ranked by WT compartment score and split into deciles (A to B). Gene expression of genes within each decile (1 to 10) was plotted for KO and WT. Number of significantly different genes between KO and WT that fell within each decile indicated. Boxes represent median and interquartile range, whiskers represent 1.5xIQR, with outliers plotted as individual points. E: Comparison of gene expression within compartment deciles between KO and WT. Boxes represent median and interquartile range, whiskers represent 1.5xIQR, with outliers plotted as individual points. Adjusted P-values for all comparisons (KO vs WT) > 0.9.

## REFERENCES

1. Fyodorov, D.V., Zhou, B.-R., Skoultchi, A.I. and Bai, Y. (2018) Emerging roles of linker histones in regulating chromatin structure and function. Nature Reviews Molecular Cell Biology, 19, 192–206.

2. Lennox, R.W. and Cohen, L.H. (1983) The histone H1 complements of dividing and nondividing cells of the mouse. Journal of Biological Chemistry, 258, 262–268.

3. Wang, Z.F., Sirotkin, A.M., Buchold, G.M., Skoultchi, A.I. and Marzluff, W.F. (1997) The mouse histone H1 genes: gene organization and differential regulation. J Mol Biol, 271, 124–138.

4. Popova, E.Y., Grigoryev, S.A., Fan, Y., Skoultchi, A.I., Zhang, S.S. and Barnstable, C.J. (2013) Developmentally Regulated Linker Histone H1c Promotes Heterochromatin Condensation and Mediates Structural Integrity of Rod Photoreceptors in Mouse Retina*. Journal of Biological Chemistry, 288, 17895–17907.

5. PiÑA, B., MartÍNez, P. and Suau, P. (1987) Changes in H1 complement in differentiating rat-brain cortical neurons. European Journal of Biochemistry, 164, 71–76.

6. Sirotkin, A.M., Edelmann, W., Cheng, G., Klein-Szanto, A., Kucherlapati, R. and Skoultchi, A.I. (1995) Mice develop normally without the H1(0) linker histone. Proc Natl Acad Sci U S A, 92, 6434–6438.

7. Fan, Y., Sirotkin, A., Russell, R.G., Ayala, J. and Skoultchi, A.I. (2001) Individual somatic H1 subtypes are dispensable for mouse development even in mice lacking the H1(0) replacement subtype. Mol Cell Biol, 21, 7933–7943.

8. Fan, Y., Nikitina, T., Morin-Kensicki Elizabeth, M., Zhao, J., Magnuson Terry, R., Woodcock Christopher, L. and Skoultchi Arthur, I. (2003) H1 Linker Histones Are Essential for Mouse Development and Affect Nucleosome Spacing In Vivo. Molecular and Cellular Biology, 23, 4559–4572.

9. Yusufova, N., Kloetgen, A., Teater, M., Osunsade, A., Camarillo, J.M., Chin, C.R., Doane, A.S., Venters, B.J., Portillo-Ledesma, S., Conway, J. et al. (2021) Histone H1 loss drives lymphoma by disrupting 3D chromatin architecture. Nature, 589, 299–305.

10. Tafon-Brown, K., Loveday, C., Yost, S., Clarke, M., Ramsay, E., Zachariou, A., Elliof, A., Wylie, H., Ardissone, A., Rignger, O. et al. (2017) Mutations in Epigenetic Regulation Genes Are a Major Cause of Overgrowth with Intellectual Disability. Am J Hum Genet, 100, 725–736.

11. Burkardt, D.D., Zachariou, A., Loveday, C., Allen, C.L., Amor, D.J., Ardissone, A., Banka, S., Bourgois, A., Coubes, C., Cytrynbaum, C. et al. (2019) HIST1H1E heterozygous protein-truncating variants cause a recognizable syndrome with intellectual disability and distinctive facial gestalt: A study to clarify the HIST1H1E syndrome phenotype in 30 individuals. Am J Med Genet A, 179, 2049–2055.

12. Takenouchi, T., Uehara, T., Kosaki, K. and Mizuno, S. (2018) Growth pafern of Rahman syndrome. American Journal of Medical GeneDcs Part A, 176, 712–714.

13. Pelle, A., Pezzoli, L., Apuril, E., Iascone, M. and Selicorni, A. (2021) A novel HIST1HE pathogenic variant in a girl with macrocephaly and intellectual disability: a new case and review of literature. Clinical dysmorphology, 30, 39–43.

14. Tremblay, M.W., Green, M.V., Goldstein, B.M., Aldridge, A.I., Rosenfeld, J.A., Streff, H., Tan, W.D., Craigen, W., Bekheirnia, N., Al Tala, S., et al. (2022) Mutations of the histone linker H1–4 in neurodevelopmental disorders and functional characterization of neurons expressing C-terminus frameshik mutant H1.4. Human Molecular GeneDcs, 31, 1430–1442.

15. Suzuki, K., Tsunekawa, Y., Hernandez-Benitez, R., Wu, J., Zhu, J., Kim, E.J., Hatanaka, F., Yamamoto, M., Araoka, T., Li, Z. et al. (2016) In vivo genome editing via CRISPR/Cas9 mediated homology-independent targeted integration. Nature, 540, 144–149.

16. George, S.H.L., Gertsenstein, M., Vintersten, K., Korets-Smith, E., Murphy, J., Stevens, M.E., Haigh, J.J. and Nagy, A. (2007) Developmental and adult phenotyping directly from mutant embryonic stem cells. Proceedings of the NaDonal Academy of Sciences, 104, 4455–4460.

17. Borghesani, P.R., Peyrin, J.M., Klein, R., Rubin, J., Carter, A.R., Schwartz, P.M., Luster, A., Corfas, G. and Segal, R.A. (2002) BDNF stimulates migration of cerebellar granule cells. Development, 129, 1435–1442.

18. Bilimoria, P.M. and Bonni, A. (2008) Cultures of Cerebellar Granule Neurons. Cold Spring Harbor Protocols, 2008, pdb.prot5107.

19. Wei, X., Xiang, Y., Peters, D.T., Marius, C., Sun, T., Shan, R., Ou, J., Lin, X., Yue, F., Li, W. et al. (2022) HiCAR is a robust and sensitive method to analyze open-chromatin-associated genome organization. Molecular Cell, 82, 1225–1238.e1226.

20. Wei, X., Tran, D. and Diao, Y. (2023) HiCAR: Analysis of Open Chromatin Associated Long-range Chromatin Interaction Using Low-Input Materials. Current Protocols, 3, e899.

21. Lyons, S.M., Cunningham, C.H., Welch, J.D., Groh, B., Guo, A.Y., Wei, B., Whinield, M.L., Xiong, Y. and Marzluff, W.F. (2016) A subset of replication-dependent histone mRNAs are expressed as polyadenylated RNAs in terminally differentiated tissues. Nucleic Acids Research, 44, 9190–9205.

22. Marzluff, W.F., Wagner, E.J. and Duronio, R.J. (2008) Metabolism and regulation of canonical histone mRNAs: life without a poly(A) tail. Nature Reviews GeneDcs, 9, 843–854.

23. Ito-Ishida, A., Yamalanchili, H.K., Shao, Y., Baker, S.A., Heckman, L.D., Lavery, L.A., Kim, J.Y., Lombardi, L.M., Sun, Y., Liu, Z. et al. (2018) Genome-wide distribution of linker histone H1.0 is independent of MeCP2. Nat Neurosci, 21, 794–798.

24. Cao, K., Lailler, N., Zhang, Y., Kumar, A., Uppal, K., Liu, Z., Lee, E.K., Wu, H., Medrzycki, M., Pan, C. et al. (2013) High-Resolution Mapping of H1 Linker Histone Variants in Embryonic Stem Cells. PLOS GeneDcs, 9, e1003417.

25. Watson, J.A., Alexander-Howden, B.K., Hall, T.S., Wear, M.A., McGhie, F., Clifford, G., Wapenaar, H., Zou, J., Bird, A. and Wilson, M.D. (2026) MeCP2 requires interactions with nucleosome linker DNA to read chromatin DNA methylation. Nature CommunicaDons, 17, 5374.

26. Ghosh, R.P., Horowitz-Scherer, R.A., Nikitina, T., Shlyakhtenko, L.S. and Woodcock, C.L. (2010) MeCP2 binds cooperatively to its substrate and competes with histone H1 for chromatin binding sites. Mol Cell Biol, 30, 4656–4670.

27. Skene, P.J., Illingworth, R.S., Webb, S., Kerr, A.R.W., James, K.D., Turner, D.J., Andrews, R. and Bird, A.P. (2010) Neuronal MeCP2 Is Expressed at Near Histone-Octamer Levels and Globally Alters the Chromatin State. Molecular Cell, 37, 457–468.

28. Salinas-Pena, M., Serna-Pujol, N. and Jordan, A. (2024) Genomic profiling of six human somatic histone H1 variants denotes that H1X accumulates at recently incorporated transposable elements. Nucleic Acids Research, 52, 1793–1813.

29. Nielsen, A.L., Oulad-Abdelghani, M., Ortiz, J.A., Remboutsika, E., Chambon, P. and Losson, R. (2001) Heterochromatin Formation in Mammalian Cells: Interaction between Histones and HP1 Proteins. Molecular Cell, 7, 729–739.

30. Serna-Pujol, N., Salinas-Pena, M., Mugianesi, F., Le Dily, F., Marti-Renom, M.A. and Jordan, A. (2022) Coordinated changes in gene expression, H1 variant distribution and genome 3D conformation in response to H1 depletion. Nucleic Acids Research, 50, 3892–3910.

31. Frank, C.L., Liu, F., Wijayatunge, R., Song, L., Biegler, M.T., Yang, M.G., Vockley, C.M., Safi, A., Gersbach, C.A., Crawford, G.E., et al. (2015) Regulation of chromatin accessibility and Zic binding at enhancers in the developing cerebellum. Nat Neurosci, 18, 647–656.

32. Tan, L., Shi, J., Moghadami, S., Parasar, B., Wright, C.P., Seo, Y., Vallejo, K., Cobos, I., Duncan, L., Chen, R. et al. (2023) Lifelong restructuring of 3D genome architecture in cerebellar granule cells. Science, 381, 1112–1119.

33. Fan, Y. and Skoultchi, A.I. (2004) Genetic analysis of H1 linker histone subtypes and their functions in mice. Methods Enzymol, 377, 85–107.

34. Marzluff, W.F. (2005) Metazoan replication-dependent histone mRNAs: a distinct set of RNA polymerase II transcripts. Current Opinion in Cell Biology, 17, 274–280.

35. Fornasiero, E.F., Mandad, S., Wildhagen, H., Alevra, M., Rammner, B., Keihani, S., Opazo, F., Urban, I., Ischebeck, T., Sakib, M.S. et al. (2018) Precisely measured protein lifetimes in the mouse brain reveal differences across tissues and subcellular fractions. Nature CommunicaDons, 9, 4230.

36. Hergeth, S.P., Dundr, M., Tropberger, P., Zee, B.M., Garcia, B.A., Daujat, S. and Schneider, R. (2011) Isoform-specific phosphorylation of human linker histone H1.4 in mitosis by the kinase Aurora B. Journal of Cell Science, 124, 1623–1628.

37. Roque, A., Ponte, I., Arrondo, J.L.R. and Suau, P. (2008) Phosphorylation of the carboxy-terminal domain of histone H1: effects on secondary structure and DNA condensation. Nucleic Acids Research, 36, 4719–4726.

38. Talasz, H., Helliger, W., Puschendorf, B. and Lindner, H. (1996) In Vivo Phosphorylation of Histone H1 Variants during the Cell Cycle. Biochemistry, 35, 1761–1767.

39. Wiśniewski, J.R., Zougman, A., Krüger, S. and Mann, M. (2007) Mass Spectrometric Mapping of Linker Histone H1 Variants Reveals Multiple Acetylations, Methylations, and Phosphorylation as Well as Differences between Cell Culture and Tissue * S. Molecular & Cellular Proteomics, 6, 72–87.

40. Joseph, F.M. and Young, N.L. (2023) Histone variant-specific post-translational modifications. Seminars in Cell & Developmental Biology, 135, 73–84.

41. Chen, Y., Hoover, M.E., Dang, X., Shomo, A.A., Guan, X., Marshall, A.G., Freitas, M.A. and Young, N.L. (2016) Quantitative Mass Spectrometry Reveals that Intact Histone H1 Phosphorylations are Variant Specific and Exhibit Single Molecule Hierarchical Dependence *. Molecular & Cellular Proteomics, 15, 818–833.

42. Chu, C.-S., Hsu, P.-H., Lo, P.-W., Scheer, E., Tora, L., Tsai, H.-J., Tsai, M.-D. and Juan, L.-J. (2011) Protein Kinase A-mediated Serine 35 Phosphorylation Dissociates Histone H1.4 from Mitotic Chromosome *. Journal of Biological Chemistry, 286, 35843–35851.

43. Saha, A., Seward, C.H., Stubbs, L. and Mizzen, C.A. (2020) Site-Specific Phosphorylation of Histone H1.4 Is Associated with Transcription Activation. InternaDonal Journal of Molecular Sciences, 21, 8861.

44. Liao, R. and Mizzen, C.A. (2017) Site-specific regulation of histone H1 phosphorylation in pluripotent cell differentiation. EpigeneDcs & ChromaDn, 10, 29.

45. Zheng, Y., John, S., Pesavento, J.J., Schultz-Norton, J.R., Schiltz, R.L., Baek, S., Nardulli, A.M., Hager, G.L., Kelleher, N.L. and Mizzen, C.A. (2010) Histone H1 phosphorylation is associated with transcription by RNA polymerases I and II. Journal of Cell Biology, 189, 407–415.

46. Prendergast, L. and Reinberg, D. (2021) The missing linker: emerging trends for H1 variant-specific functions. Genes Dev, 35, 40–58.

47. Willcockson, M.A., Healton, S.E., Weiss, C.N., Bartholdy, B.A., Botbol, Y., Mishra, L.N., Sidhwani, D.S., Wilson, T.J., Pinto, H.B., Maron, M.I. et al. (2021) H1 histones control the epigenetic landscape by local chromatin compaction. Nature, 589, 293–298.

48. Lister, R., Mukamel, E.A., Nery, J.R., Urich, M., Puddifoot, C.A., Johnson, N.D., Lucero, J., Huang, Y., Dwork, A.J., Schultz, M.D. et al. (2013) Global Epigenomic Reconfiguration During Mammalian Brain Development. Science, 341, 1237905.

49. Clowney, E.J., LeGros, Mark A., Mosley, Colleen P., Clowney, Fiona G., Markenskoff-Papadimitriou, Eirene C., Myllys, M., Barnea, G., Larabell, Carolyn A. and Lomvardas, S. (2012) Nuclear Aggregation of Olfactory Receptor Genes Governs Their Monogenic Expression. Cell, 151, 724–737.

50. Solovei, I., Kreysing, M., Lanctôt, C., Kösem, S., Peichl, L., Cremer, T., Guck, J. and Joffe, B. (2009) Nuclear Architecture of Rod Photoreceptor Cells Adapts to Vision in Mammalian Evolution. Cell, 137, 356–368.

