## supplemental figures for "The linker histone H1.4 condenses chromatin in maturing postmitotic neurons"

s1

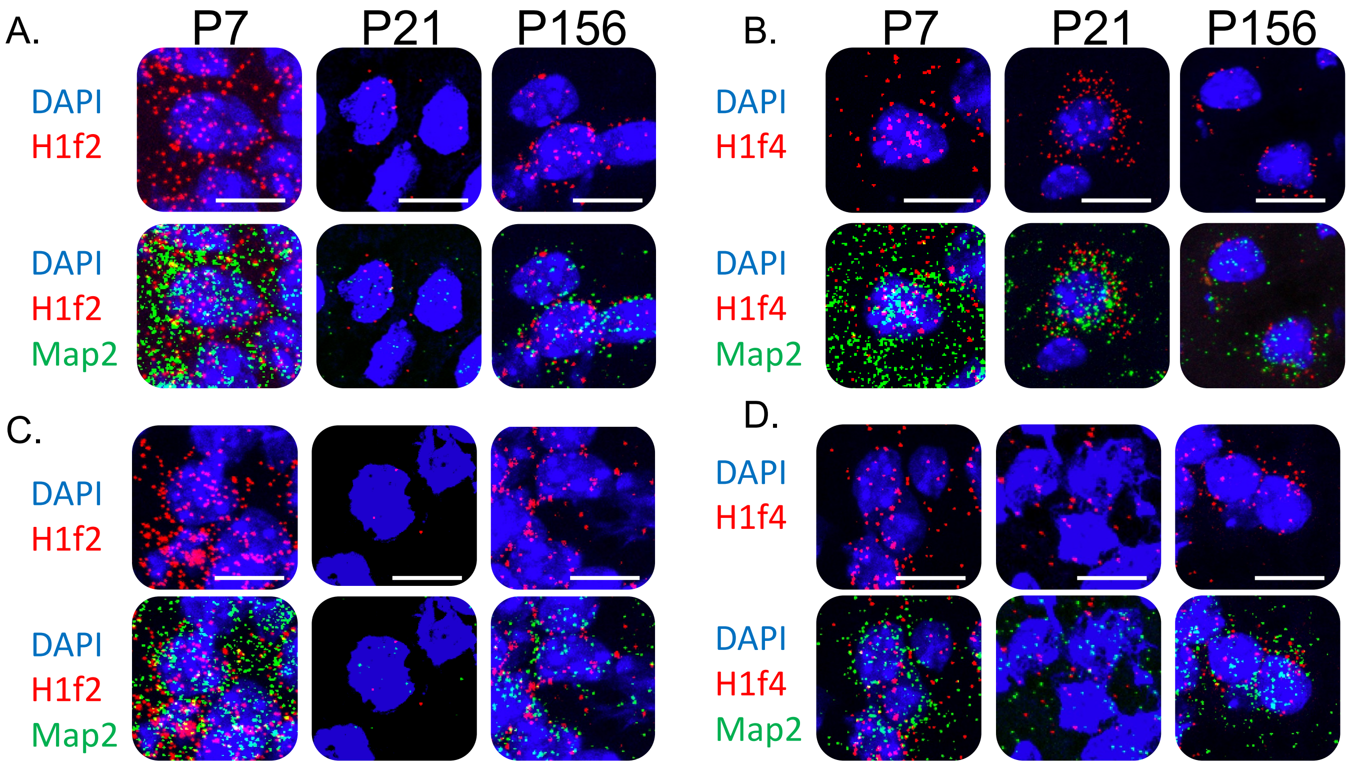
A.
P7
P21
P156
DAPI
H1f2
DAPI
H1f2
Map2
B.
P7
P21
P156
DAPI
H1f4
DAPI
H1f4
Map2
C.
DAPI
H1f2
DAPI
H1f2
Map2
D.
DAPI
H1f4
DAPI
H1f4
Map2

S2

A.

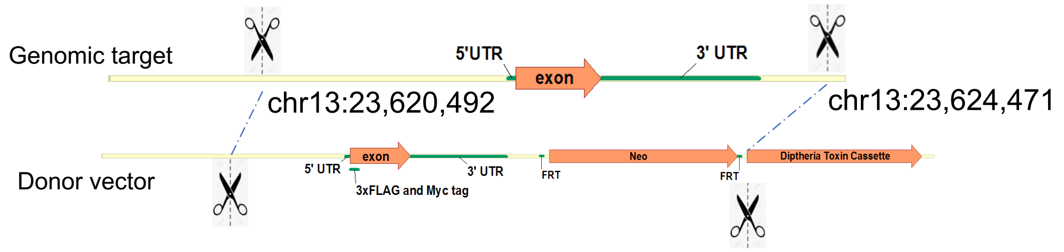
Genomic target
5'UTR
exon
3' UTR
chr13:23,620,492
chr13:23,624,471
Donor vector
5' UTR
exon
3' UTR
3xFLAG and Myc tag
FRT
Neo
FRT
Diptheria Toxin Cassette

B.

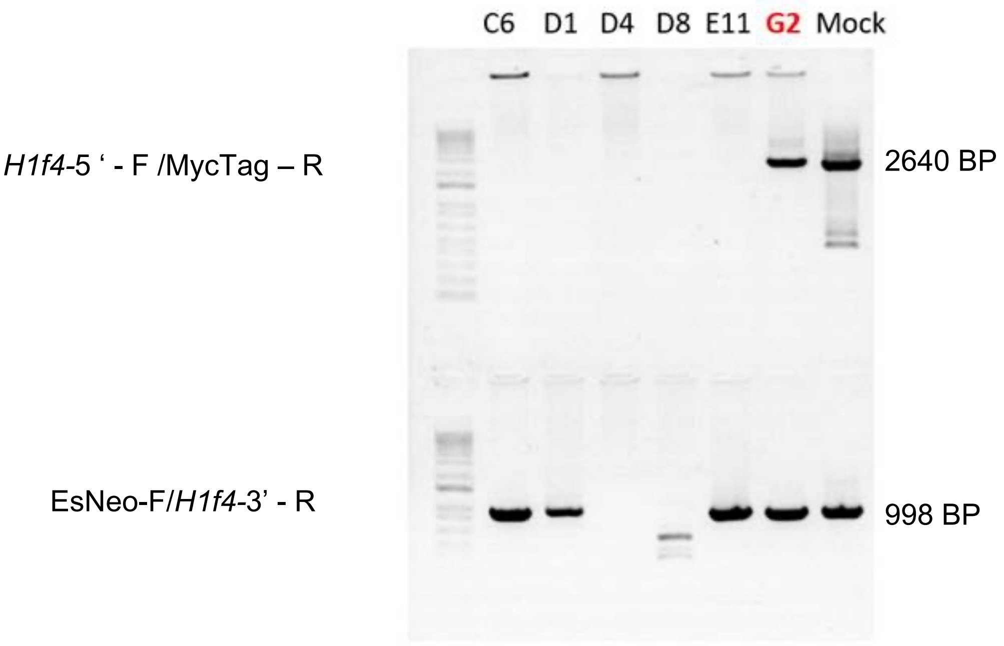
C6 D1 D4 D8 E11 G2 Mock
*H1f4*-5 ' - F /MycTag – R
2640 BP
EsNeo-F/*H1f4*-3' - R
998 BP

C.

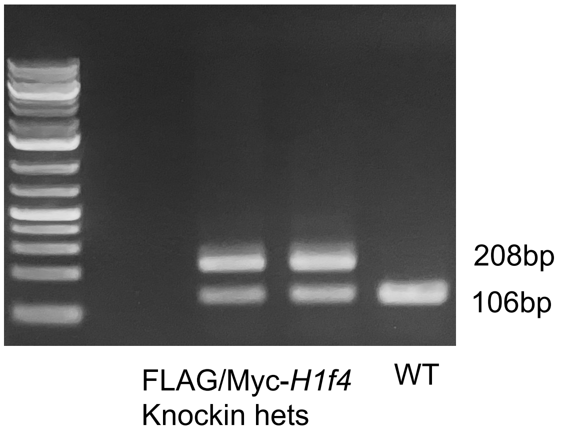
208bp
106bp
FLAG/Myc-*H1f4*
Knockin hets
WT

S3

Cerebellum

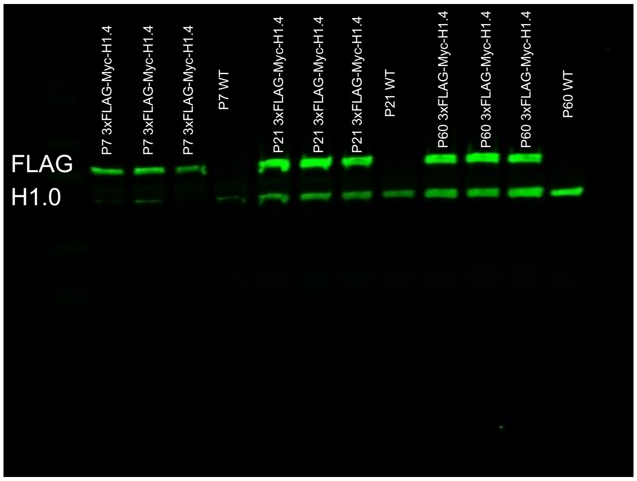


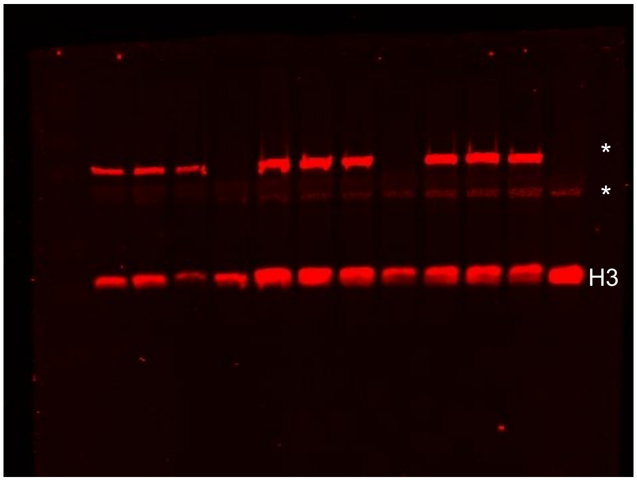


*Bleed through from other channel

Cortex

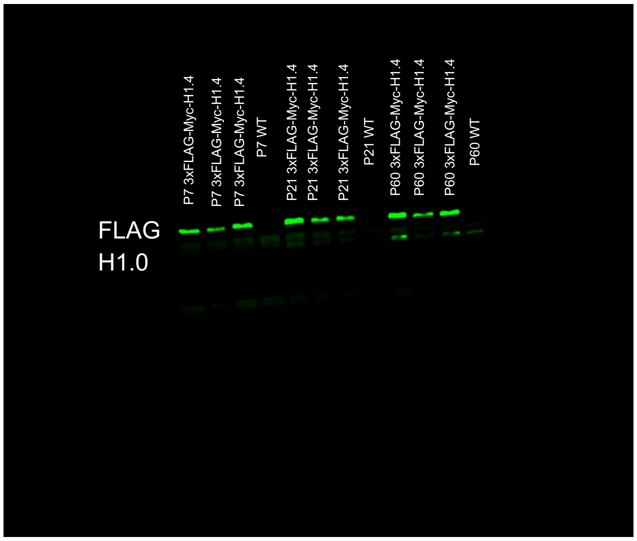


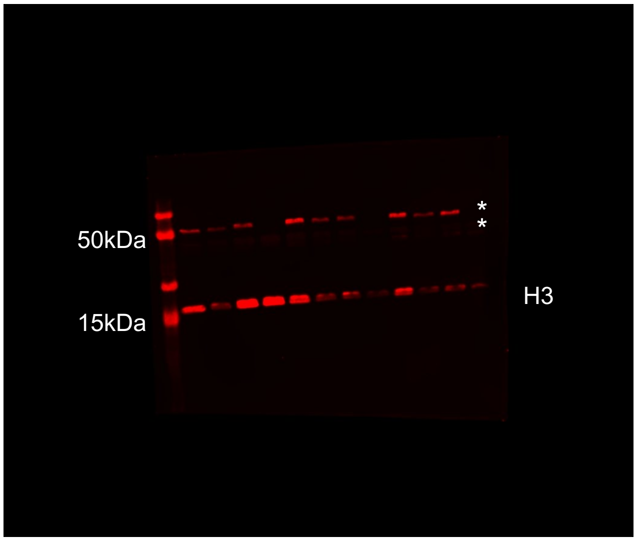


*Bleed through from other channel

Hippocampus

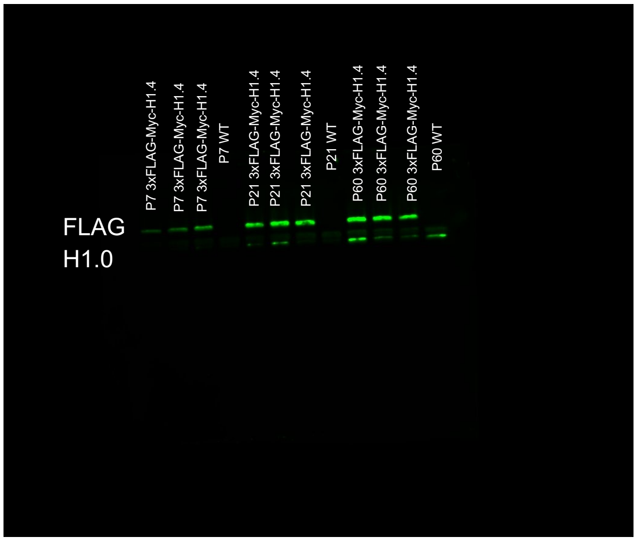


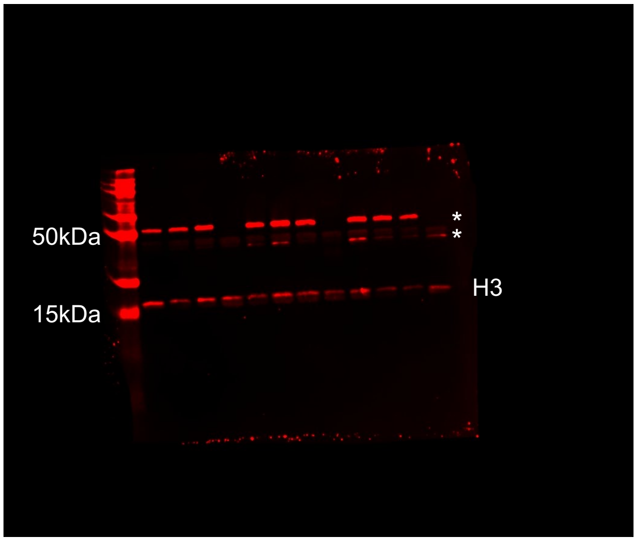


*Bleed through from other channel

S4

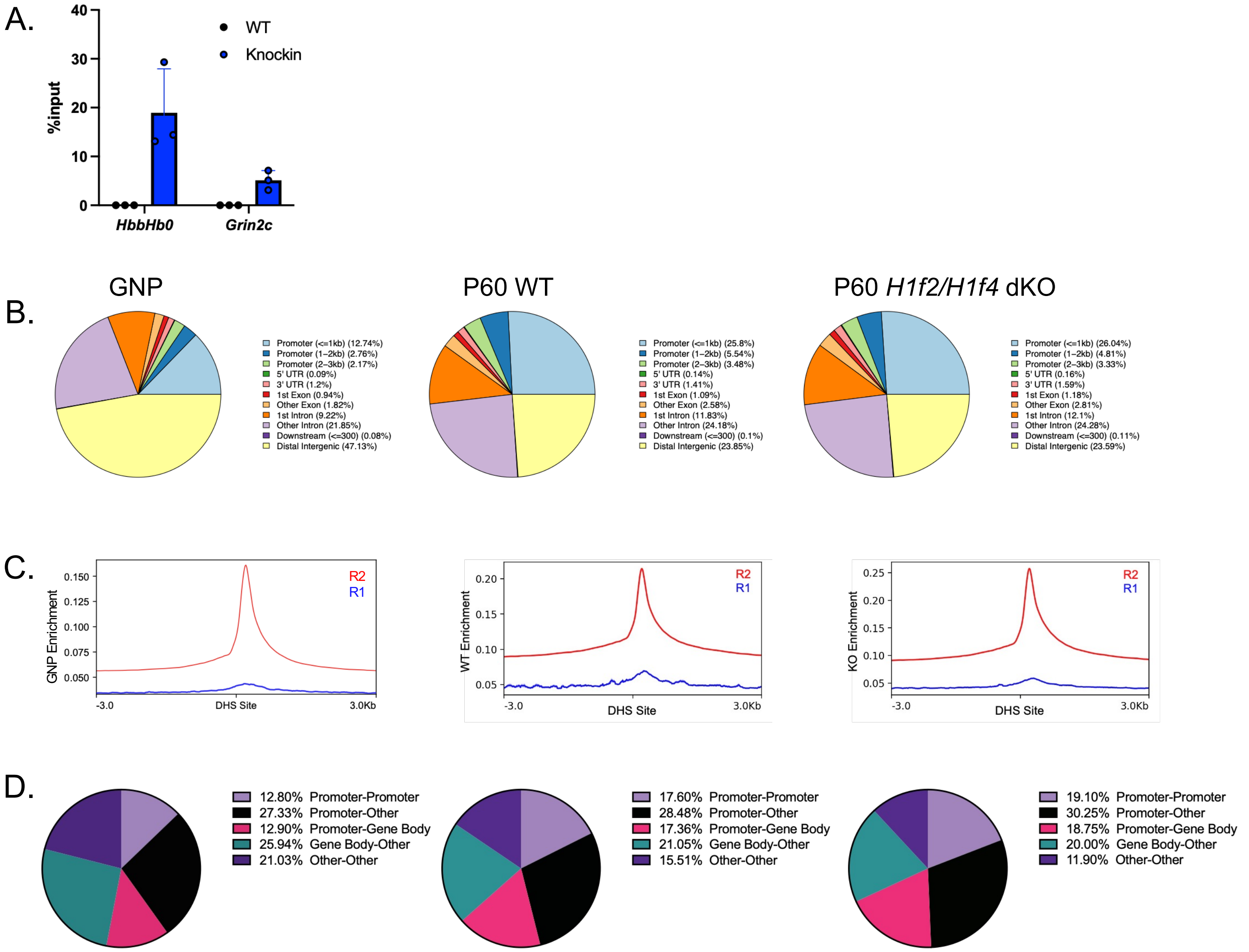
A.
B.
C.
D.
WT
Knockin
%input
40
30
20
10
0
HbbHb0
Grin2c
GNP
P60 WT
P60 H1f2/H1f4 dKO
Promoter (<=1kb) (12.74%)
Promoter (1–2kb) (2.76%)
Promoter (2–3kb) (2.17%)
5' UTR (0.09%)
3' UTR (1.2%)
1st Exon (0.94%)
Other Exon (1.82%)
1st Intron (9.22%)
Other Intron (21.85%)
Downstream (<=300) (0.08%)
Distal Intergenic (47.13%)
Promoter (<=1kb) (25.8%)
Promoter (1–2kb) (5.54%)
Promoter (2–3kb) (3.48%)
5' UTR (0.14%)
3' UTR (1.41%)
1st Exon (1.09%)
Other Exon (2.58%)
1st Intron (11.83%)
Other Intron (24.18%)
Downstream (<=300) (0.1%)
Distal Intergenic (23.85%)
Promoter (<=1kb) (26.04%)
Promoter (1–2kb) (4.81%)
Promoter (2–3kb) (3.33%)
5' UTR (0.16%)
3' UTR (1.59%)
1st Exon (1.18%)
Other Exon (2.81%)
1st Intron (12.1%)
Other Intron (24.28%)
Downstream (<=300) (0.11%)
Distal Intergenic (23.59%)
GNP Enrichment
0.150
0.125
0.100
0.075
0.050
R2
R1
-3.0
DHS Site
3.0Kb
WT Enrichment
0.20
0.15
0.10
0.05
KO Enrichment
0.25
12.80% Promoter-Promoter
27.33% Promoter-Other
12.90% Promoter-Gene Body
25.94% Gene Body-Other
21.03% Other-Other
17.60% Promoter-Promoter
28.48% Promoter-Other
17.36% Promoter-Gene Body
21.05% Gene Body-Other
15.51% Other-Other
19.10% Promoter-Promoter
30.25% Promoter-Other
18.75% Promoter-Gene Body
20.00% Gene Body-Other
11.90% Other-Other

s5

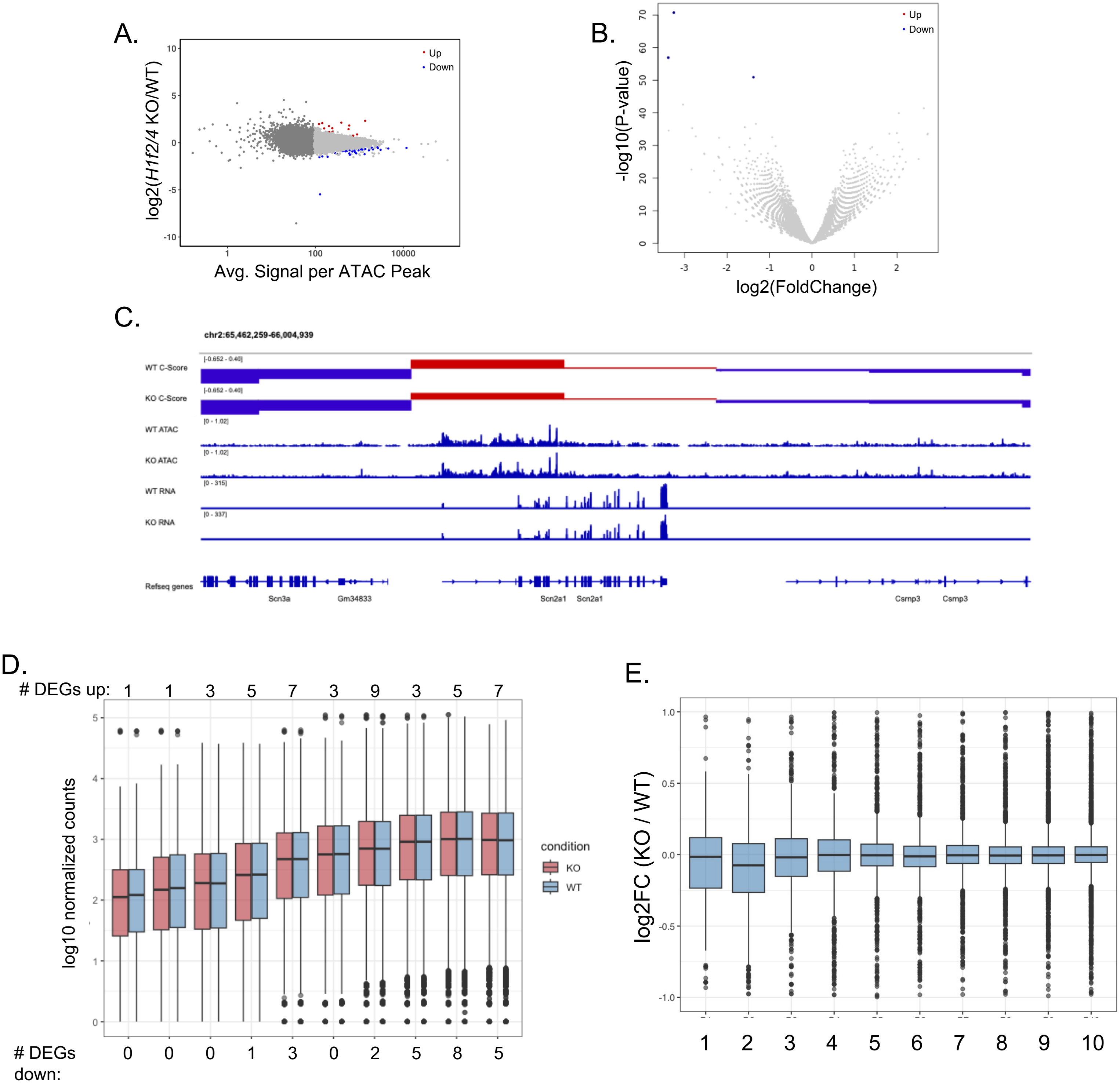
A.
log2(H1f2/4 KO/WT)
Avg. Signal per ATAC Peak
Up
Down
B.
-log10(P-value)
log2(FoldChange)
Up
Down
C.
chr2:65,462,259-66,004,939
WT C-Score
KO C-Score
WT ATAC
KO ATAC
WT RNA
KO RNA
Refseq genes
Scn3a
Gm34833
Scn2a1
Scn2a1
Csmp3
Csmp3
D.
### DEGs up: 1 1 3 5 7 3 9 3 5 7
log10 normalized counts
condition
KO
WT
### DEGs down: 0 0 0 1 3 0 2 5 8 5
E.
log2FC (KO / WT)
1 2 3 4 5 6 7 8 9 10
